# TipA, a *Bdellovibrio bacteriovorus* BPI-like double-TULIP is opened by its adapter protein, TipB

**DOI:** 10.1101/2025.09.09.675085

**Authors:** Simon G. Caulton, Ian T. Cadby, Gareth W. Hughes, Hannah Johnston, Timothy J. Knowles, Andrew L. Lovering

## Abstract

*Bdellovibrio bacteriovorus* is a predatory bacterium that invades the periplasm of other Gram-negative bacteria and liberates prey biomolecules for replication. *Bdellovibrio* has a wealth of genes that encode unique proteins to enable this lifestyle. Using a series of x-ray structures, we show that the operonal pairing of *bd2538* and *bd2539,* encode a double <u>T</u>ULIP (tubular lipid binding protein) invasion <u>p</u>rotein A (TipA) and a small beta sandwich (TipB), respectively. TipA has a specialised N-terminal TULIP domain with a beta hairpin and helix that mediate homodimerisation through hairpin interdigitation. This dimerisation creates a large, continuous, enclosed lumen that we demonstrate to contain multiple lipid molecules. In addition, we show that TipB functions as a small adapter protein that binds to TipA using specialised loops that bury into the TipA hydrophobic core. This binding forms a clamp on the edge of the N-terminal beta sheet and induces a large 20 Å conformational change, opening the TULIP fold to create an accessible interior. This study presents the first structural characterisation of lipid binding proteins in *Bdellovibrio*, and the first example of conformational change in TULIPs mediated by an adaptor protein.

**Synopsis:** 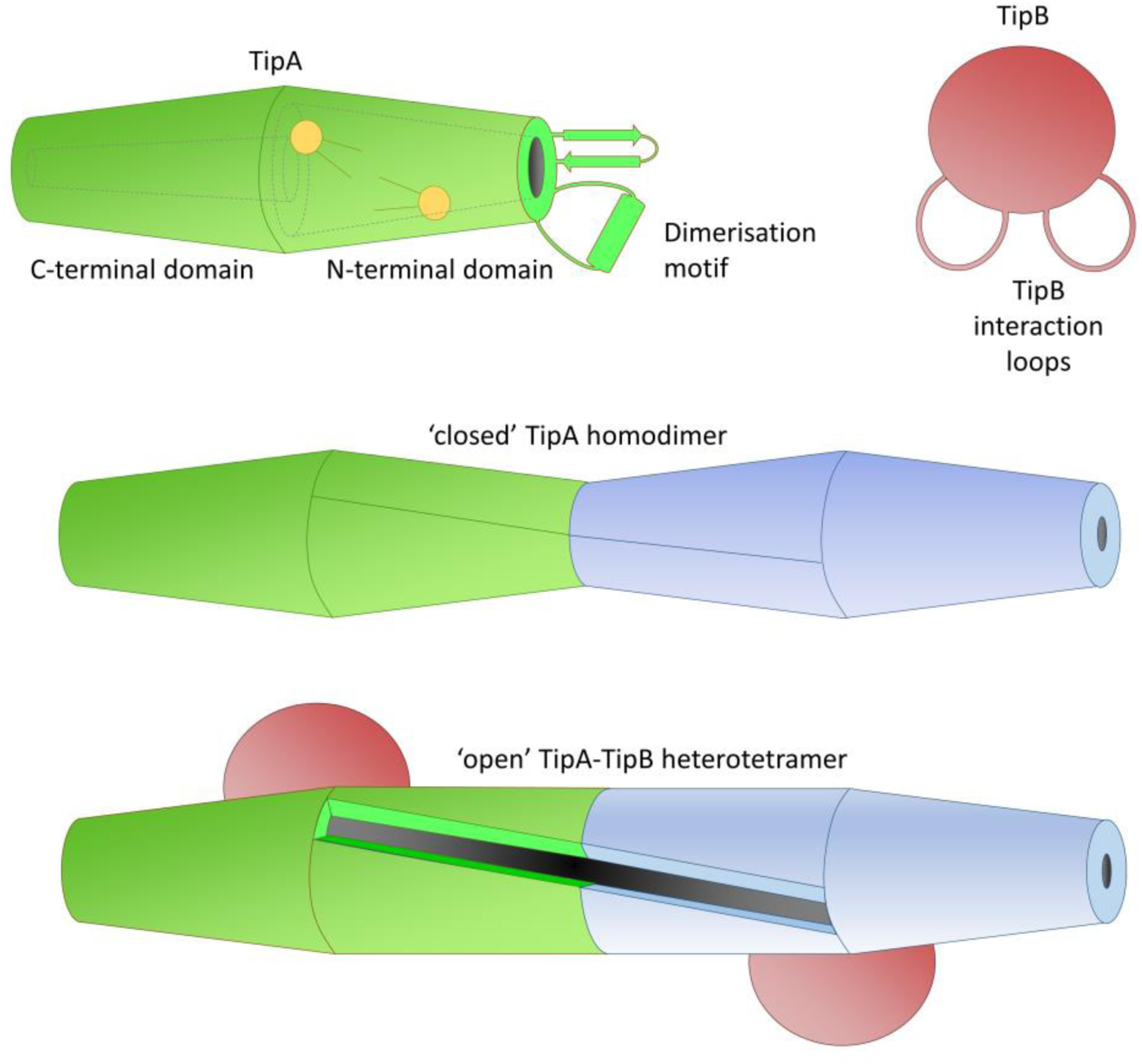

*Bdellovibrio bacteriovorus* is a predatory bacterium that invades the periplasm of other Gram-negative bacteria in order to consume them from within. To enable this lifestyle, it has an arsenal of predation-associated genes, including the operonal pair *bd2538* and *bd2539*. We use structural and biophysical techniques to characterise two proteins produced by these genes, TipA and TipB.

- TipA is a double tubular lipid binding protein (TULIP) that forms a homodimer that sequesters lipids in its lumen and binds to lipid bilayers
- TipB is a small beta sandwich that interacts with TipA via two specialised loops
- Interaction of TipA with TipB opens the TULIP fold of TipA

## Introduction

*Bdellovibrio bacteriovorus* is a predatory bacterium that preys on other Gram-negative bacteria. Upon prey encounter, *B. bacteriovorus* attaches to its prey and invades its periplasm, where it remodels the host peptidoglycan to form a rounded bdelloplast. Here, it liberates prey biomolecules for its filamentous growth, followed by septation into progeny. Finally, when the prey cell is exhausted of nutrients, it becomes lysed to release free swimming *B. bacteriovorus* to repeat the cycle anew [1]. This remarkable prey-killing lifestyle, and its indifference to prey antibiotic-resistance status gives *Bdellovibrio* huge potential as a novel biological control agent [2]. However, this lifestyle also requires a wealth of specialised tools [3]. Indeed, *Bdellovibrio* utilizes a number of liberated biomolecules from its prey [4–7], including both lipid A [8] and fatty acids [9]. These lipids can either remain in their original form or undergo metabolic alterations by *Bdellovibrio*.

*Bdellovibrio*’s intricate association with membranes is central to its predatory lifestyle. Membrane and cell wall remodelling [10–12], as well as lipid uptake [8, 9] are critical facets of host cell penetration and subsequent intraperiplasmic survival. The mechanisms through which *Bdellovibrio* obtains lipids from its prey remain elusive, highlighting the complexity of lipid transport in bacteria, which typically necessitates dedicated machinery, such as the Lpt or Mla systems for lipid movement across the aqueous periplasmic environment [13]. However, these systems are responsible for intracellular transport of lipids across the periplasm of a cell, shuttling between inner and outer membranes. *Bdellovibrio* has the additional need for intercellular lipid transport, to extract and transfer lipids from prey cell membranes to self.

Tubular lipid-binding proteins (TULIPs) exhibit unique structural features comprising a cylindrical fold that surrounds a hydrophobic cavity, enabling them to sequester lipids from the surrounding environment. These include the extracellular bacterial permeability increasing protein-like (BPI-like) double-TULIP family, Takeout-like proteins and intracellular Synaptotagmin-like mitochondrial-lipid-binding protein (SMP) family [14, 15]. TULIPs have gained attention for their diverse roles in membrane tethering, lipid transfer, hormone binding and innate immunity in eukaryotes [15, 16]. Recently, widespread identification in prokaryotes has made it clear that TULIPs have important roles in prokaryote physiology [17, 18]. Structurally characterised prokaryotic TULIPS are few, and include *E. coli* YceB, and *Clostridium botulinum* neurotoxins/neurotoxin-associated proteins, P47 and OrfX proteins [19, 20]. Understanding the molecular mechanisms governing TULIP-lipid interactions is crucial for deciphering their role in prokaryotic organisms, inclusive of alternative lifestyles like the predator *Bdellovibrio*.

In this study, we provide X-ray crystal structures of two proteins encoded from the operonal gene cluster *bd2538* and *bd2539,* TULIP invasion protein A (TipA) and TULIP-associated invasion protein B (TipB), respectively. These proteins are upregulated during the invasion process [21] and secreted into prey [22]. We show that TipA forms a novel dimeric BPI-like double-TULIP, longer than classical characterized family representatives, and that TipB binds to TipA to induce a conformational opening of the TULIP fold. These results provide the first example of a lipid-binding protein in *Bdellovibrio* and provide insight into the molecular mechanisms of lipid trafficking during its predatory lifestyle.

## Results

### Bdellovibrio TipA is a BPI-like double TULIP divergent from other family members

The operonal genes *bd2538* and *bd2539* (Fig. 1A) encode two proteins we term TULIP invasion protein A (TipA) and TULIP-associated invasion protein B (TipB), respectively. To identify the role of TipA in predation we sought to determine its structure (mature enzyme, minus signal peptide) and elucidate its function. The crystal structure of selenomethionine-incorporated TipA was solved in space group P 3_1_ 2 at 4 Å resolution with three molecules in the asymmetric unit (ASU). The low resolution of diffraction was likely a consequence of the 87% solvent content caused by the end-to-end packing of the proteins in the unit cell. Nevertheless, maps of sufficient quality were obtained (Fig. 1B) and in all three chains, at least residues N35-A566 were traced with no unmodelled gaps in the model (Fig. 1C). TipA was annotated to contain a DUF2785 domain, however the crystal structure confirms mis-annotation, as the protein fold shows it to be of the lipid binding protein BPI/LBP superfamily (IPR030675) [23].

**Figure 1.**
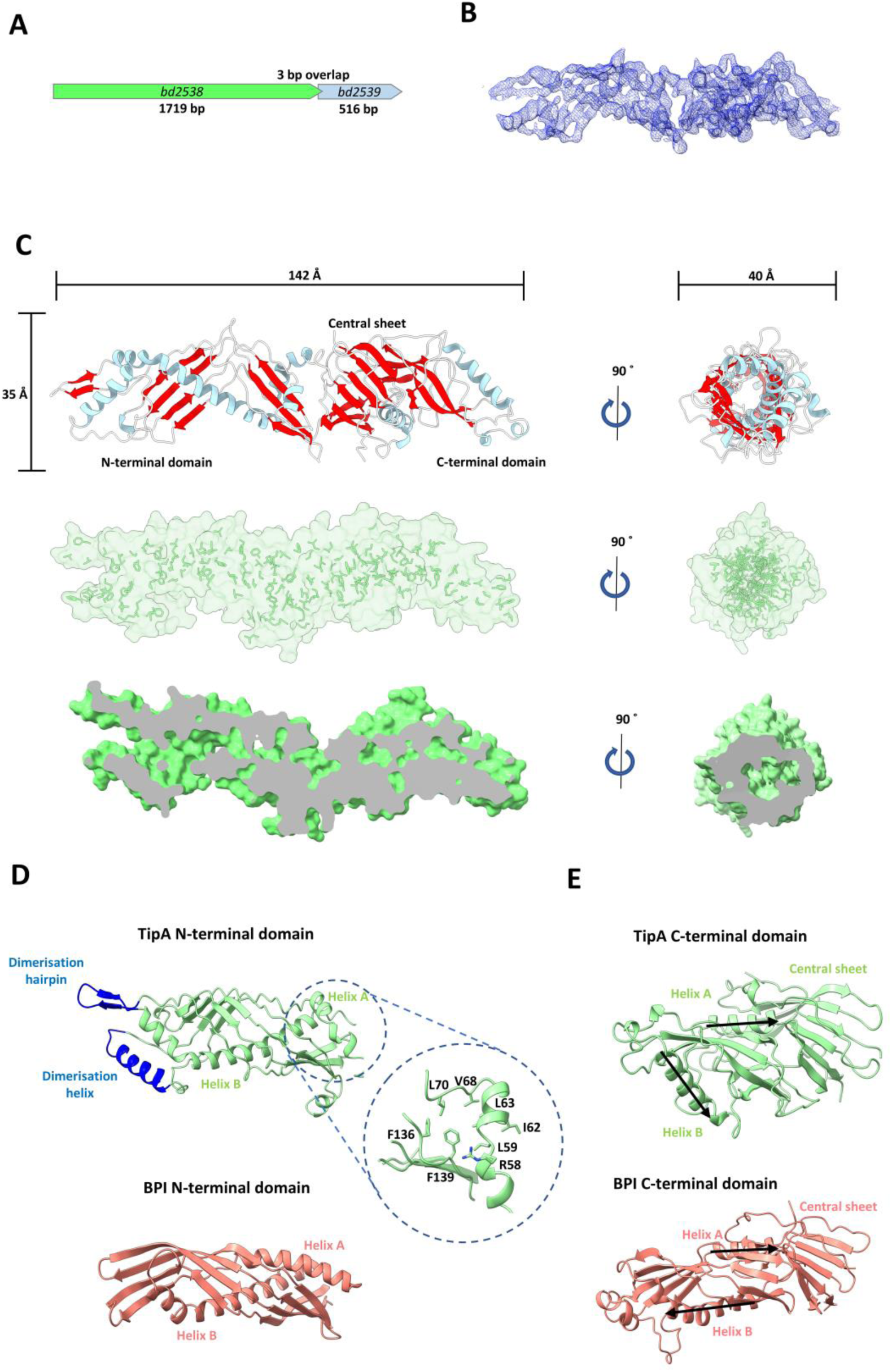
The structure of *Bdellovibrio* TULIP protein TipA. **(A)** Schematic of the forward strand ORFs for the gene operon of *bd2538* (green) and *bd2539* (blue). **(B)** 2Fo-Fc map at 2.5 σ surrounding chain C of the crystal structure of TipA after the final refinement. **(C)** Top; Crystal structure of TipA, showing the double-TULIP fold of the Bactericidal permeability-increasing protein, alpha/beta domain superfamily, coloured by secondary structure (beta – red, helix – light blue, coil – white). The lengths of the protein are shown for all three dimensions. Middle; a surface view of TipA with hydrophobic residues shown in stick form, highlighting the hydrophobic nature of the central cavity. Bottom; A clipped surface view showing the inner cavity. (**D)** Top; structural features of the N-terminal TipA TULIP domain, with the dimerisation motif coloured blue. The expanded view shows residues in stick form that cause the N-terminal Helix A to protrude. Bottom; The structure of the N-terminal TULIP domain of bacterial permeability increasing protein (BPI, code 1BP1) for comparison. **(E)** Top; structural features of the C-terminal TipA TULIP domain, with the helix positions highlighted with arrows. The expanded view shows residues in stick form that cause the N-terminal Helix A to protrude. Bottom; The structure of the C-terminal TULIP domain of BPI for comparison.

TipA shows similar gross features to those of other BPI/LBP family structures: a double-TULIP fold, comprising a central beta sheet sandwiched by two tubular domains each formed by a beta sheet that wraps around an extended helix (Fig. 1C). Each TULIP domain has the typical secondary structure characteristic of that domain: an N-terminal helix (helix A), an elongated beta sheet, and a C-terminal helix (helix B). The single protomer forms a cylindrical protein with dimensions of approximately 142 x 35 x 40 Å enveloping a central hydrophobic cavity throughout the entire protein.

TipA was submitted to the DALI server [24] to identify proteins with similar structures. The top hits were all double TULIP proteins: Bacterial permeability-increasing protein (BPI; Z-score 16.5; 21% seq ID), Cholesteryl ester transfer protein (CETP; Z-scores 13.5-15) and LPS-binding protein (LBP; Z-scores 10.4-11). Superposition of the individual N- and C-terminal domains of TipA with those of BPI shows significant divergence of the protein fold, with RMSD values of 5.7 Å over 176 residues for the N-terminal domain and 5.8 Å over 112 residues for the C-terminal domain (Fig. 1D-E). Hence, we conclude that TipA is a divergent TULIP representing a grouping of the superfamily previously uncharacterised.

The N-terminal domain shows three inserts relative to that of BPI: a beta hairpin between the last beta strand and helix B, a loop containing an additional helix between the first and second strands, and a small helix between the second and third strands (Fig. 1D). The first two of these inserts comprise the dimerisation interface of TipA, whereas the function of the latter is unclear.

Compared to BPI, the C-terminal domain has a truncated loop between strands 1 and 2, a larger triangular-shaped loop between strands 2 and 3, and two small helices inserted between the last strand and helix B. Superposition reveals that the central beta sheet and helix A align well with the BPI C-terminal domain, but upon exit from helix A, TipA diverges greatly (Fig. 1E). In particular, both the sheet and helix B are much shorter than BPI, and this causes a relative displacement.

### TipA forms a cylindrical homodimer

A surprising finding of the crystal structure of TipA was that unlike any other characterised TULIP, TipA forms a C2 symmetrical dimer (Fig. 2A), with one complete dimer in the ASU and one protein forming a dimer with a symmetry copy. TipA has a novel beta hairpin and helix on the N-terminal domain that form a dimer interface, producing an extended hydrophobic tube through an end-on-end interaction (Fig. 2A-B). The channel throughout dimeric TipA is almost continuous, containing a pinch point in each of the central sheet domains, and has a total volume of approximately 25000 Å^3^ as calculated by 3vee [25] (Fig. 2B). The dimerisation hairpins form a single antiparallel beta sheet and produce hydrophobic and polar faces with two intermolecular electrostatic interactions formed by E251 and K256 (Fig. 2C). A sequence alignment of TipA shows a number of highly conserved residues that are important structurally (Fig. EV1). The tip of the dimerisation hairpin contains a highly conserved asparagine (N253) that hydrogen bonds with the backbone of F184. N260 is also highly conserved and serves to propagate helix B following the hairpin. The highly conserved P246 kinks the backbone to bend the hairpin away from the beta sheet and angle it parallel to the protein tube, allowing interdigitation with the hairpin of the second molecule. The N-terminus of the dimerisation helix packs against its own helix B and extends outwards to pack its C-terminal tryptophan into the hydrophobic face of the other protomer’s helix B (Fig. 2D). The dimerisation helix makes hydrogen bonds between Q104 and Q104, and E263 and Q111. The associated loop (L113-S117) forms backbone hydrophobic interactions through L113 and F116 and backbone hydrogen bonds between V114 and Y258. A highly conserved glycine (G115) allows the chain to form a sharp turn to thread back to the beta sheet of the N-terminal domain.

**Figure 2.**
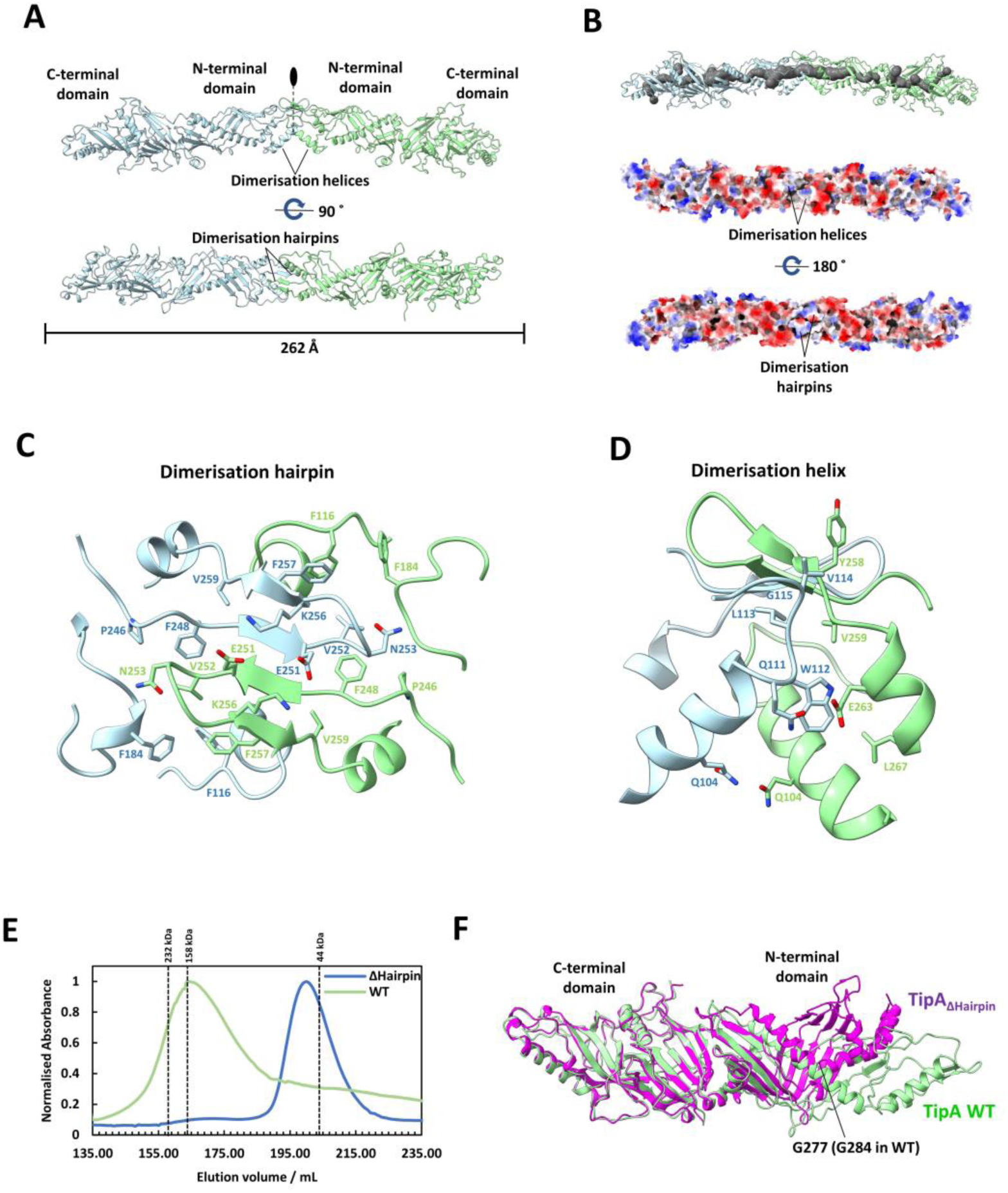
Dimerisation of TipA into a four TULIP-long assembly. **(A)** Crystal structure of TipA, showing two chains forming an end-on-end homodimer via interaction of their N-terminal TULIP domains. One protomer is shown in light blue and the other is in green with the 2-fold symmetry axis shown as a black oval and dotted line. **(B)** Upper; the dimer forms an elongated cavity within (grey surface, calculated by Mole [52]). Lower; the electrostatic surface view shows that the dimer forms a continuous tube with very few solvent accessible gaps. **(C)** The view down the two-fold axis shows how the two interdigitated dimerisation hairpins (light blue and green) create a polar surface on the outside of the TULIP dimer and a hydrophobic inner face for a continuous hydrophobic channel. **(D)** The dimerisation helix forms polar and hydrophobic interactions with the other protomer’s Helix B and dimerisation helix. **(E)** Mutation of the dimerisation hairpin causes a shift in the elution volume of TipA on gel filtration from 165 mL to 200 mL. **(F)** The crystal structure of TipA_ΔHairpin_ (magenta) superimposed onto the TipA wild type (WT) structure (green). The C-terminal domain overlays well, but the N-terminal domain is contracted due to twisting of the beta-sheet and bending of the N-terminal Helix B at G277.

Due to the spacious packing of the TipA dimer crystals, and consequent relatively low resolution, we subsequently sought to obtain a higher resolution structure of TipA by altering the packing of the crystal lattice. To do this we substituted the hairpin region (A249-F258) with a triple glycine linker, termed TipA_ΔHairpin_. We expected the loss of dimerisation to allow better packing of the protein, reducing solvent content to produce higher resolution diffraction. This substitution shifted the elution volume of the protein from 164 mL to 200 mL on a Superdex 200 pg 26/60 column, showing a drastic decrease in hydrodynamic radius of the protein, and indicating that the protein is primarily monomeric in solution (Fig. 2E). As predicted, the TipA_ΔHairpin_ protein crystallised with different packing in space group P 2_1_ 2_1_ 2_1_, with a solvent content of 60% and higher resolution diffraction of 2.6 Å. The crystal structure of TipA_ΔHairpin_ contains a single monomeric copy in the ASU (Fig. 2F). The lack of dimer interface introduced disorder into the N-terminal domain and the regions of G107-M118 and the glycine linker region P246-N253 (N260 in wild-type (WT)) so these residues could not be modelled. The C-terminal domain and central sheet are very similar to that of the WT protein, but the N-terminal TULIP sheet is pulled inward, mediated by a bend in Helix B at G277 (G284 in WT). This suggests that dimerisation is integral to maintaining correct folding of the N-terminal domain.

### TULIP partner TipB is a β-sandwich with exposed hydrophobic loops

Since the TipA-encoding *bd2538* gene is in a predicted operon with the downstream *bd2539* gene, which it overlaps by 4 nucleotides and which we name *tipB*, we hypothesised that TipB could be functionally linked to TipA. Following our structural studies of TipA, we purified and crystalized the cryptic operonal TipB protein. TipB crystallised in space group C 2 2 2_1_ with three copies in the ASU and diffracted to 2.17 Å resolution. The structure reveals that TipB forms a compact β-sandwich, in which one four-strand anti-parallel sheet forms the central globular core of the protein and is packed on one side by a pair of short beta sheets and the other side by an elongated capping-loop extending from β7 and β8 (Fig. 3A). This loop folds back on the sheet burying Y120, W122 and M133 into the core. The protein has another pair of loops that extend between β3 and β4, and β6 and β7, termed interaction loops 1 and 2, respectively (Fig. 3A). Importantly, both loops have hydrophobic residues at the ends: F65 and I66 in loop 1 and F103 in loop 2, and demonstrate a high degree of flexibility (as inferred by electron density and B-factor values; Fig 3B). Finally, the protein contains a short helical turn between strands 2 and 3 and an 11-residue helix between strands 8 and 9.

**Figure 3.**
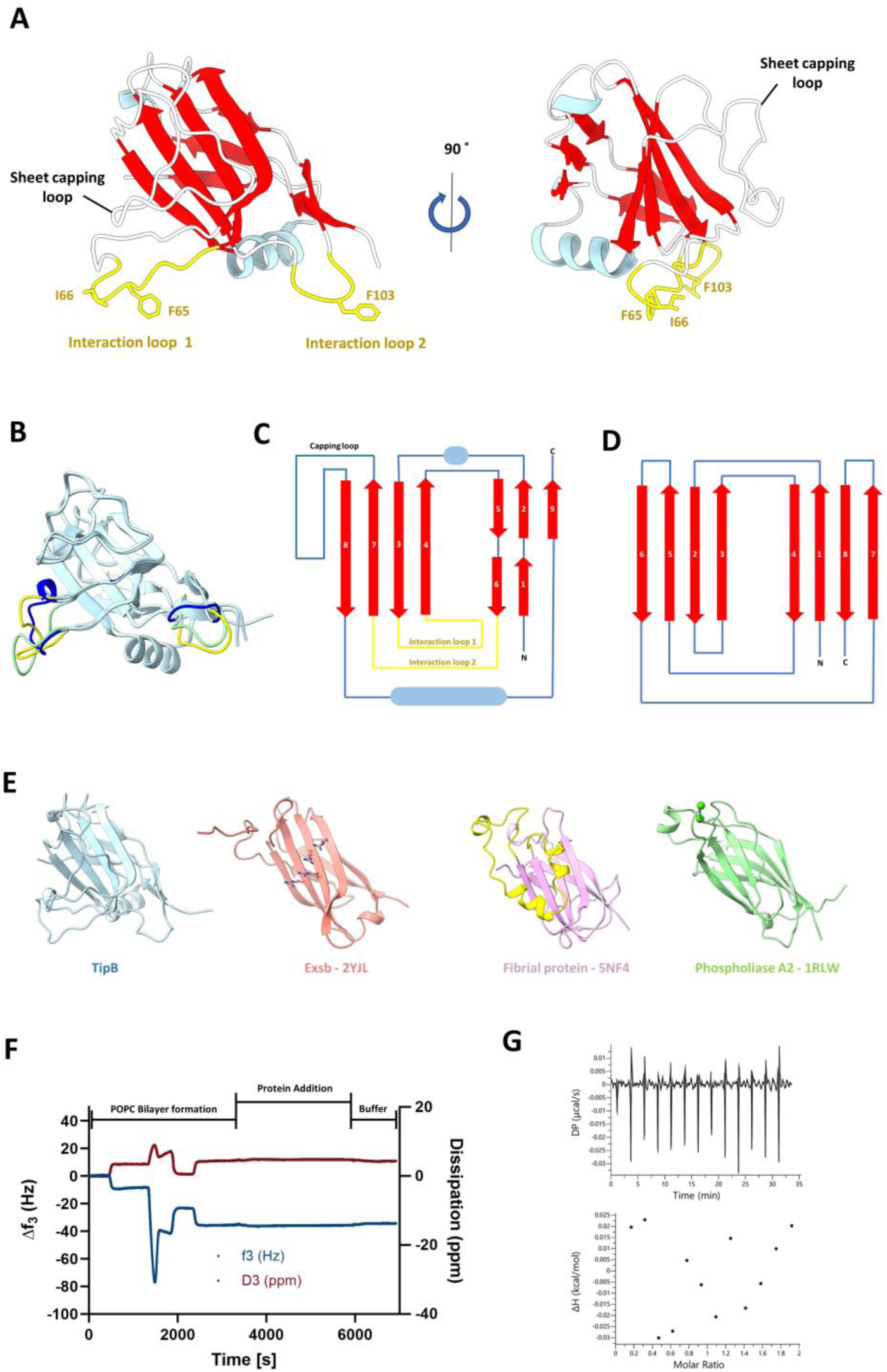
Crystal structure of the TipB adapter protein. **(A)** Crystal structure of TipB, coloured by secondary structure (beta – red, helix – light blue, coil – white) with the two specialised interaction loops coloured yellow. Key hydrophobic amino acids in the interaction loops are shown in stick form. **(B)** The three chains of TipB in the ASU are superposed to highlight flexibility in the interaction loops (shown blue, green and yellow for each chain). **(C)** The topology of TipB, with strands in red, helices as blue ovals and loops as blue lines. **(D)** The topology of type II C2 domains. **(E)** Crystal structures of proteins identified in structural homology searches with DALI [24], shown in same orientation as TipB (left, light blue). Arginines that form the basic face of the sheet in *Pseudomonas aeruginosa* ExsB [26] (coloured salmon) are shown in stick form; the sheet capping loop of *Porphyromonas gingivalis* [53] (coloured pink) is shown in yellow; and the two calcium ions of human phospholipase A2 [35] (coloured green) are shown as spheres. **(F)** QCM-D trace showing f3 and d3 signals. Upon protein addition, no significant change in f3 or d3 is observed. **(G)** ITC thermogram of a titration of calcium chloride into TipB showing no binding.

We used the DALI server [24] to identify structural homologues of TipB and the highest scoring structure was *P. aeruginosa* ExsB [26] (PDB code 2YJL), with a Z-score of 7.3. Both TipB and ExsB show a topology similar to that of type II C2 domains (Fig. 3C-E) [27], a calcium-dependent membrane-localisation domain found in multiple eukaryotic signalling proteins including kinases [28], lipases [29] and membrane trafficking proteins [30]. In comparison, TipB lacks the 8^th^ strand that runs antiparallel between the 1^st^ and 7^th^ strands, instead packing its last strand parallel to the 2^nd^ strand (the 1^st^ strand is broken into two smaller strands due to a broken beta sheet in TipB), forming a mixed sheet. The other top-scoring hits all had the standard type II C2-domain topology, with varying additional loops and secondary structure components. Most of the C2-domain proteins have an exposed face on the beta sheet, however some homologues show extensions in the β7 and β8 loop that cap off this sheet in a similar manner to TipB (Fig. 3E). None of the homologues had extensions between β3 and β4 or β6 and β7 strands where the interaction loops are observed in TipB.

ExsB has an exposed beta sheet with a basic face predicted to mediate binding to acidic lipids (Fig. 3E). In TipB, this sheet face is not exposed and forms part of the hydrophobic core that is enclosed by an extended loop between β7 and β8. To investigate whether TipB would adhere to a lipid bilayer, we used quartz crystal microbalance with dissipation monitoring (QCM-D) measurements (Fig. 3F). This surface-based technique monitors changes to the resonant frequency of a surface (Δf) under an applied voltage. Δf is inversely proportional to the mass deposited on the surface, and as such allows us to monitor interactions between biomolecules [31]. In the case of TipB-lipid interactions, a stable POPC bilayer was deposited onto a silicon QCM-D sensor, following well-established methods [32, 33]. Despite not being a constituent of bacterial membranes, a POPC bilayer was used here due to its well-reported ability to form highly stable bilayers when deposited onto a surface [34]. Once lipid bilayers were deposited, TipB was introduced and its interaction with the lipid bilayer was assessed in a semi-quantitative manner. Upon flow of TipB over the bilayer, negligible frequency (0.3 Hz) and dissipation (0.48 ppm) changes occurred, thus confirming that TipB alone does not bind directly to lipid membranes.

C2 domains commonly bind calcium, such as the high-scoring DALI hit *Homo sapiens* phospholipase 2 [35]. We wanted to confirm that TipB doesn’t bind calcium as it does not contain obvious calcium-binding residues nor was calcium observed in the crystal structure. After treatment with 5 mM EGTA and 5 mM EDTA, and subsequent gel filtration, no binding was seen using isothermal titration calorimetry (ITC; Fig. 3G), confirming that TipB is a calcium-independent C2-like β-sandwich.

### TipA and TipB form a TULIP:partner complex

Since TipB did not display any phospholipid binding activity we next hypothesised that TipA and TipB might interact with each other in a functional manner (whilst also being stable in their isolated states, as inferred from our structures). Isothermal titration calorimetry (ITC) of TipB into TipA resulted in a clear binding curve (Fig. 4A), and the -TΔS value suggests the interaction is largely driven through entropy effects, possibly through hydrophobic contributions. However, due to the low stoichiometry observed (0.25), the curve wasn’t suitable to calculate an accurate Kd. The relatively poor stability of the TipA protein would not allow for higher concentrations to be used. The low stoichiometry could result from TipA existing as multiple species in solution. Although TipA elutes as a single peak in gel filtration experiments, this peak is broad (when compared with ΔHairpin TipA), potentially the result of multiple TipA conformers (Fig. 2E). It is therefore likely that only a subpopulation of TipA is competent for binding and may further demonstrate the flexible nature of the protein.

**Figure 4.**
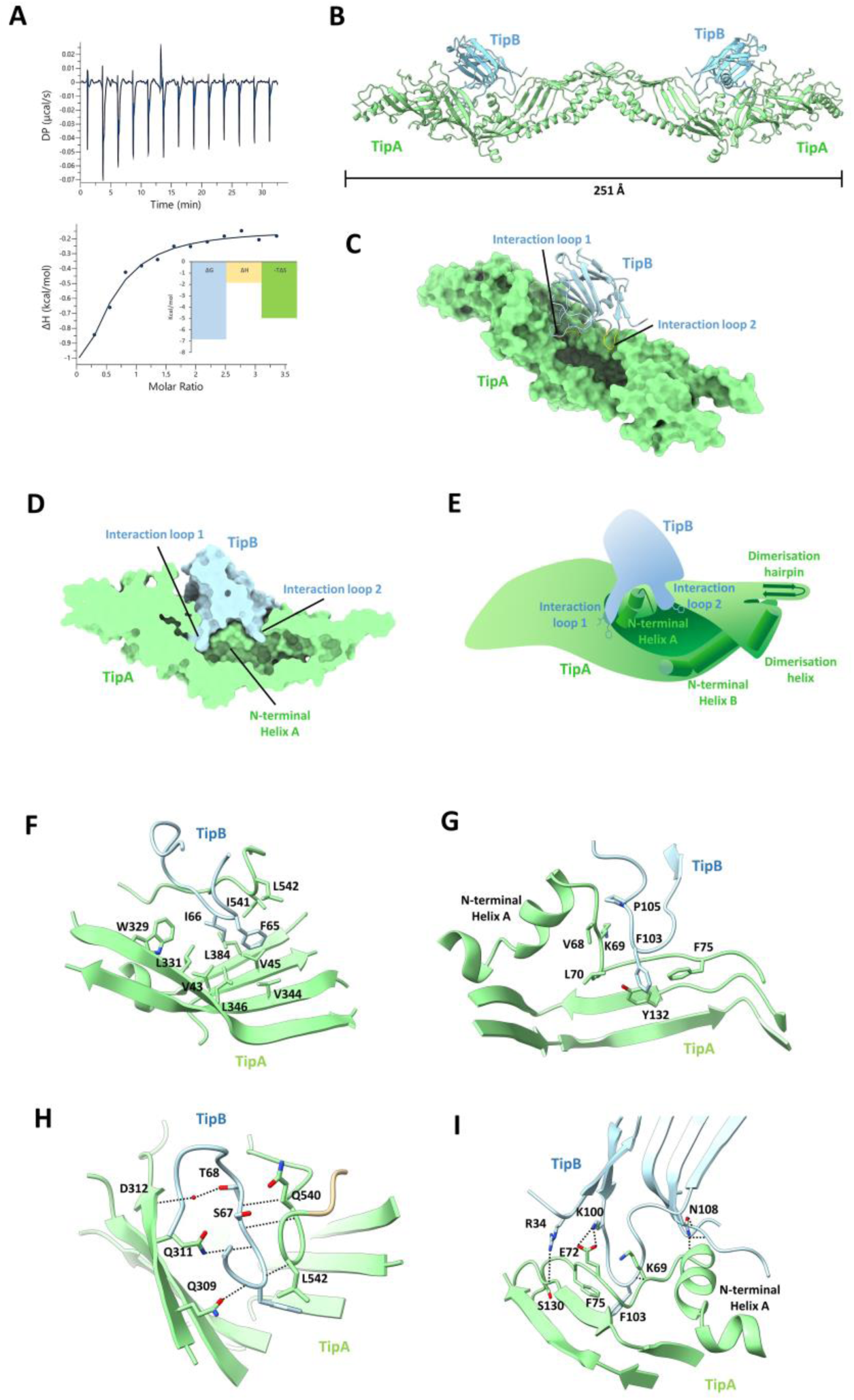
The interaction of the TipA and TipB TULIP:adapter pairing. **(A)** ITC thermogram of a titration of TipB into TipA, showing binding of the two proteins in solution. The reported energies for binding are shown as an embedded bar graph. **(B)** Crystal structure of the TipAB complex. Tip A is shown in green and forms the same 2-fold dimer as the isolated form through a crystallographic axis. TipB is shown in blue on top of TipA. **(C)** A surface view of TipA is shown, highlighting burial of the two interaction loops of TipB (coloured yellow). **(D)** A clipped surface view of the TipAB complex, showing TipA sat on the TipA N-terminal helix A. **(E)** A cartoon schematic of the interaction with parts of the TipA N-terminal TULIP domain secondary structure visible, and key hydrophobic residues of TipB shown as sticks. **(F)** Hydrophobic interactions of the TipB interaction loop 1. **(G)** Hydrophobic interactions of the TipB interaction loop 2. **(H)** polar interactions of the TipB interaction loop 1. **(I)** Polar interactions of the TipB interaction loop 2.

We subsequently desired to obtain a crystal structure of the complex of TipA and TipB. The two proteins were purified and mixed with a molar excess of TipB in the crystallisation conditions in order to ensure a complex of the two proteins. This complex crystallised in space group P 2 2 2_1_ and diffracted to 2.6 Å with one copy of each protein in the ASU. The TipA component formed a dimer on a crystallographic 2-fold axis exhibiting the same hairpin-helix dimer interface as the selenomethionine-labelled TipA crystals (Fig. 4B). TipB can be observed binding centrally on TipA, straddling the unusually kinked helix A, which protrudes away from the beta sheet, forming a triangular seat (Fig. 4C-E). The structure of complexed TipB is very similar to that of TipB alone, with RMSDs ranging from 1.43 to 1.81 Å over 139 amino acids for the three chains of the TipB crystal structure. Differences are mostly observed in interaction loops 1 and 2, which fold down over the seat and bury residues F65, I66 and F103 into the hydrophobic lumen of TipA (Fig. 4F-G). In addition to the hydrophobic components, there are a number of hydrogen and electrostatic bonds from side chains and backbone atoms throughout the 1346 Å^2^ interface (Fig. 4H-I), as calculated by PISA [36].

### Partner TipB binds to open up the TULIP fold and central cavity

The most striking observation of the complex structure is the conformational change seen in TipA’s N-terminal domain (Fig. 5A-B). While the superposition of the C-terminal domains of the apo and complexed TipA structures show an RMSD of 1.26 Å over 261 residues, the N-terminal domain shows an RMSD of 6.13 Å over 271 residues. The N-terminal domain of the TipA apo structure comprises an almost completely closed tube, with very little access to solvent (Fig. 5C). However, the N-terminal domain of the complex is in an open conformation, with a gap of approximately 20 Å (from Y80-S282; Fig. 5D). This opening is mediated by an approximate 45° rotation of the protruding TipA helix A (Fig. 5A), which acts like a lever, pulling the N-terminal sheet open. This opening is also stabilised by the anchoring of interaction loop 1 into the C-terminal domain, and the packing of interaction loop 2 F103 under the lifted N-terminal sheet. TipB F103 acts as a clamp underneath the TipA beta sheet in concert with K100 and R34, which form electrostatic interactions with E72 and E129, respectively, on the top of the sheet (Fig. 5E). Analysis by DynDom [37] characterises this as a hinge mechanism, comprising three moving domains (Fig 5F): the C-terminal domain and most of the N-terminal helix B (Domain 1); most of the N-terminal sheet and dimerisation hairpin (Domain 2); and the end of the N-terminal sheet and dimerisation helix (Domain 3). The analysis identifies two hinge points between these domains which include the rotation point of the N-terminal helix A seat. Domain 2 rotates approximately 29° relative to Domain 1 and Domain 3 rotates 36° relative to Domain 2, keeping it a similar relative position to Domain 1 in both structures (Fig. 5G). Hence, the function of TipB is to convert TipA to a more open form and, to the best of our knowledge, represents a new type of TULIP binding-partner.

**Figure 5.**
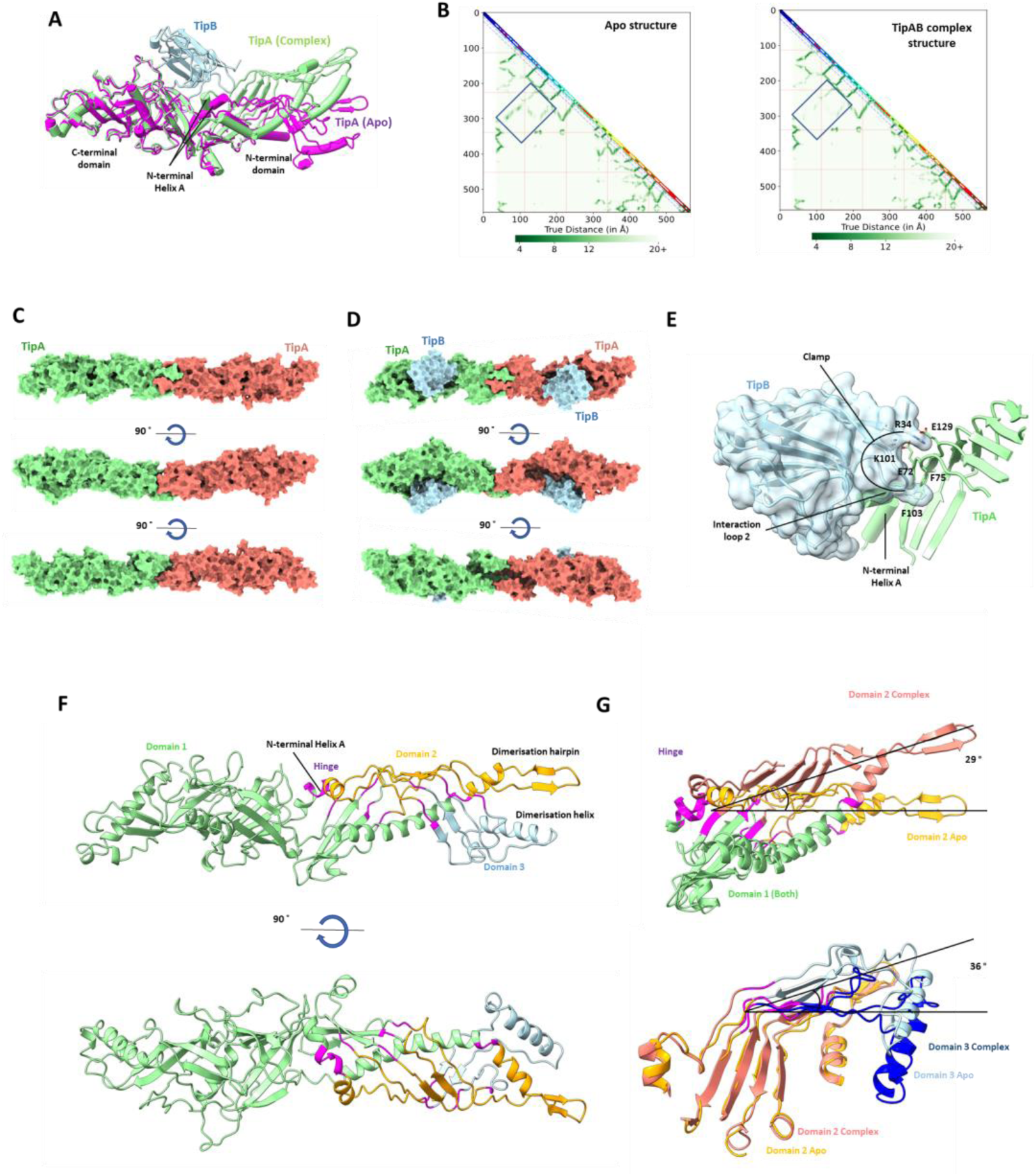
Conformational change of TipA when bound to the TipB adapter. **(A)** A superimposition of apo TipA (magenta) with the TipAB complex. Large motions in the N-terminal domain of TipA are visible. **(B)** contact plots of TipA for the apo TipA and TipAB structures, generated by DISTEVAL [54]. Both plots have a box surrounding residues that differ significantly between the two structures, which are only found in the N-terminal domain. **(C)** Three orientations of the apo TipA homodimer (one protomer green, one protomer red), showing a mostly-closed tube. **(D)** The same three orientations of the TipA homodimer as **(C)** for the TipAB complex. TipB is shown in blue, and the large opening of the TULIP homodimer stretching between the two TipB proteins is visible. **(E)** The clamp formed by interaction loop 2 and charged residues grasp the bottom and top of the TipA N-terminal beta sheet, respectively. **(F)** Analysis of the apo TipA and complexed TipA structure with DynDom to investigate the motions of the conformational change identifies three domains (coloured green, orange and blue), with motions in the hinge regions (magenta). **(G)** Upper; superpositions show the 29° rotation between Domain 1 and Domain 2 rotate by 29. Lower; superpositions show the 36° movement between domain 2 and domain 3.

Using Foldseek [38] to search for homologous proteins, we observe double-TULIPs with additional fused domains (Fig. EV2). These include beta-rich domains, helical bundles and mixed domains including enzymes and entire double-TULIP repeats. Of particular interest is the presence of a double-TULIP in the *Planctomyces sp*. SH-PL14 protein, Uniprot accession A0A142WTQ2. This protein contains a fused N-terminal esterase-like enzyme in addition to a C-terminal beta-sandwich. This beta-sandwich has similar topology to TipB, and is also predicted to localise on the N-terminal Helix A and sheet, suggesting that it could have a similar role to TipB in this system.

### TipA binds rapidly to membrane bilayer surfaces

During the purification process of TipA, we observed that under conventional buffer conditions (<500 mM NaCl) the protein remained largely in the pellet after lysis and centrifugation. However, upon addition of high salt concentrations or detergents, the protein was released into the supernatant. This led us to believe that the protein was adhering to the membrane fraction (Fig. 6A).

**Figure 6.**
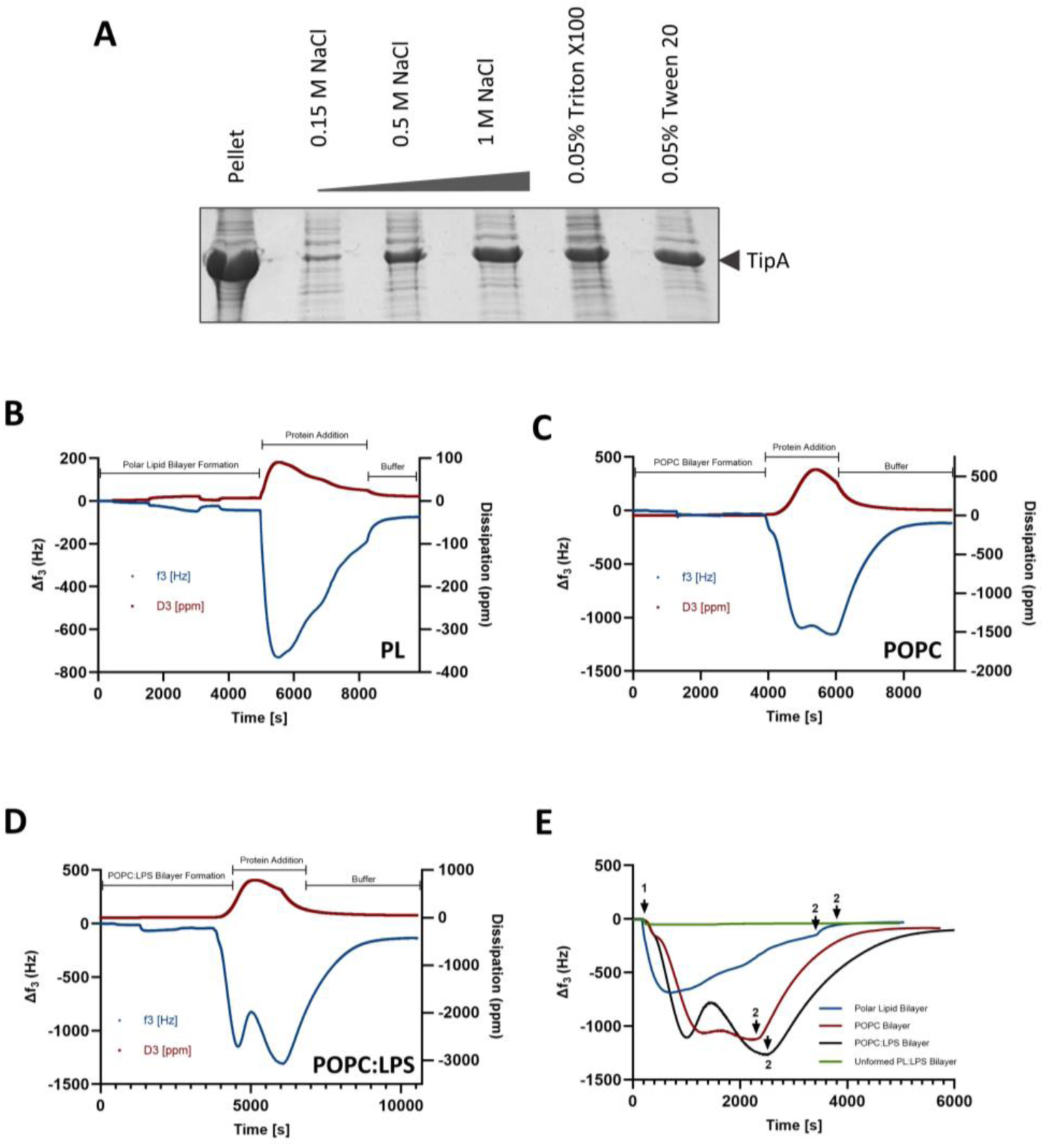
TipA can insert into a range of lipid bilayers. **(A)** Pellet-binding assays with TipA. SDS-PAGE of pellet and supernatants from various buffer washes. A prominent band for soluble TipA is observed in high salt concentrations or with the presence of detergents. **(B)** QCM-D measurements showing the changes in frequency (blue) and dissipation (red) in response to the mass deposition associated with *E. coli* polar lipid bilayer formation (polar lipid liposomes were deposited onto a silicon surface and ruptured through osmotic shock), followed by the addition of TipA, and subsequent bilayer washing with excess buffer. **(C)** QCM-D measurements as specified in **(A)** showing POPC bilayer formation, TipA addition, and buffer wash. **(D)** QCM-D measurements as specified in **(A)** indicating the formation of a stable POPC:LPS bilayer, TipA addition and subsequent buffer wash. **(E)** A comparison of the changes to frequency (indicative of mass changes), as measured through QCM-D, in response to the addition of TipA to various lipid bilayer compositions (1) and subsequent buffer wash (2). Note we were unable to successfully form a mixed polar lipid, LPS bilayer.

To confirm this, we again used QCM-D (Fig. 6B-E). To investigate TipA-lipid interactions, stable bilayers consisting of either *E. coli* polar lipids, POPC or mixed POPC:LPS lipids were deposited onto a silicon QCM-D sensor. Upon introduction of TipA to all three bilayer compositions monitored here, it exerted a dramatic negative Δf, suggesting it was rapidly deposited into the bilayer systems. This interaction was most pronounced for TipA deposition into POPC:LPS bilayers (Fig. 6D), with a Δf3 of −1106 Hz, compared to −1064 Hz and −686 Hz for POPC and polar lipid bilayers, respectively. In all bilayers tested, a critical point was reached upon which further addition of TipA resulted in removal of mass from the surface. This suggests an ability to first insert into lipid bilayers followed by membrane disruption. Furthermore, in bilayers containing POPC a double-dip insertion of TULIP was observed (Fig. 6C-D), which may result from the ability of POPC to form more stable bilayers on silicon surfaces than *E. coli* polar lipids. Interestingly, this distinctive mode of interaction was not observed when TipA was introduced to an empty silicon surface generated through the inability to deposit polar lipid:LPS bilayers, with no major mass deposition or removal observed (Fig. 6E).

### Lipids are observed in the TipA TULIP lumen in multiple states

In order to confirm that TipA is a classic TULIP that binds lipids within its cavity, thin layer chromatography (TLC) was performed (Fig. 7A). TipA clearly copurifies with all three common *E. coli* polar lipids, cardiolipin (CL), phosphatidylethanolamine (PE) and phosphatidylglycerol (PG). These have likely been extracted from *E. coli* during the expression or extraction process. Density for lipids was observed in all three crystal structures, however the low resolution of the selenomethionine form prevented their modelling (Fig. EV2A). In both the 2.6 Å TipAB complex structure (Fig. 7B) and the 2.6 Å monomeric TipA_ΔHairpin_ structure (Fig. 7C), clear densities are observed for two lipid head groups in the N-terminal TULIP domain. It was also possible to trace the lipid tails in the TipA_ΔHairpin_ structure but not in the complex structure. The lipids were modelled as PE as this headgroup fit the density well (Fig. 7D), and the density would not accommodate the glycerol moiety of PG, nor did the density indicate the presence of the double headgroup of CL.

**Figure 7.**
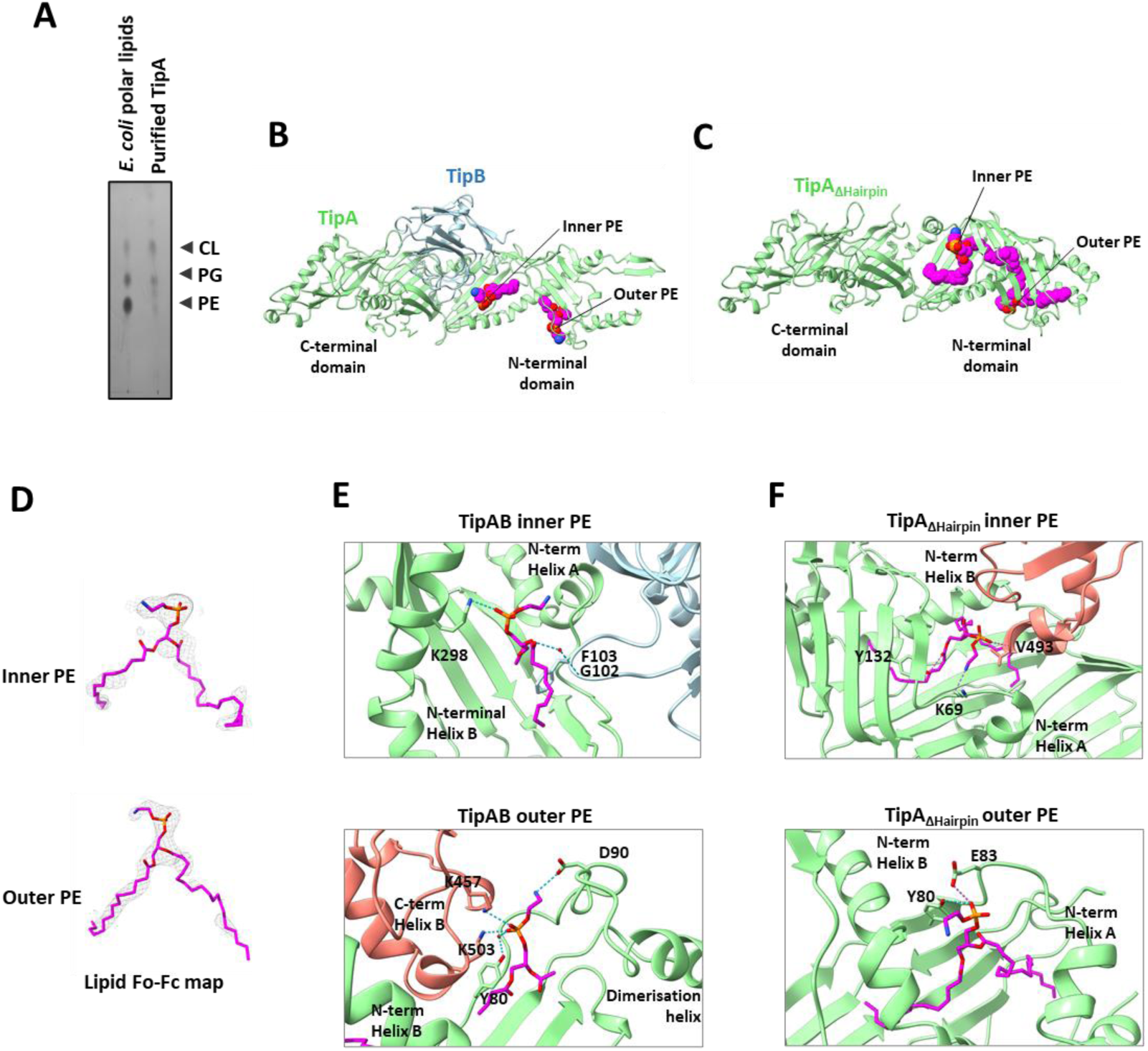
TipA sequesters lipids in the TULIP lumen. **(A)** TLC of lipids extracted from TipA. All three *E. coli* polar lipids are visible in the TipA lane (cardiolipin (CL), phosphatidylglycerol (PG), phosphatidylethanolamine (PE)), as are some additional species. **(B)** Crystal structure of the TipAB complex, with two lipid head groups shown as spheres (magenta carbons). **(C)** Crystal structure of TipAΔHairpin, with two modelled PE molecules shown as spheres (magenta carbons). **(D) 2**Fo-Fc electron density of the two lipids in the TipA_ΔHairpin_ structure at 0.8 σ. **(E)** The molecular interactions of TipAB complex and the lipid head groups for the inner PE molecule (top) and outer PE molecule (bottom). TipA is coloured green, TipB is blue, with a symmetry TipA copy coloured salmon. Lipids are shown in stick form and coloured as heteroatoms (carbon – magenta, oxygen – red, nitrogen – blue, phosphorus – orange) **(F)** The molecular interactions of TipA_ΔHairpin_ and the lipid head groups for the inner PE molecule (top) and outer PE molecule (bottom). TipA is coloured green with a symmetry TipA copy coloured salmon. Lipids are shown in stick form and coloured as heteroatoms (carbon – magenta, oxygen – red, nitrogen – blue, phosphorus – orange)

In both structures the lipid headgroups are seen in similar regions in the protein (as are the Fo-Fc peaks in the selenomethionine TipA structure; Fig. EV2A), one underneath Helix A, and the other between the sheet and Helix B (Fig. 7E-F). However, contributing amino acids that bind the ligand are different between the structures. The placement of the inner lipid in the TipAB structure means that the headgroup makes a H-bond and hydrophobic interactions with G102 and F103. In the TipA_ΔHairpin_ structure (where TipB is not present), it makes H-bonds with Y132 and K69, but additional interactions are formed by V493 of the symmetry copy, providing a backbone H-bond and sidechain hydrophobic interactions. These modelling differences suggest that TipB affects lipid interactions with TipA.

Interestingly, in both structures one of the lipids makes contact with the C-terminal domain of a symmetry copy (Fig. EV2B-D), with polar contributions from the symmetry copy K457 and K503 to the phosphate moiety of the outer lipid in the TipAB structure. When visualising higher order packing in all three TipA structures, this is a common occurrence – in three different space groups the C-terminal hydrophobic residues M454, F455, L504 and F505 pack into the N-terminal domain ‘seam’ between Helix B and the 1^st^ beta strand (Fig. EV2E-G). Further structures of close TipA homologues are needed to determine if this mode of interaction is a conserved feature.

## Discussion

Tubular lipid binding proteins (TULIP) proteins are a universal lipid-binding fold, characterised by their ability to sequester lipids within their internal hydrophobic cavities. Whilst these folds are ubiquitous to all domains of life, prokaryotic TULIPS present an under-represented group within the literature [17]. We have structurally characterised the first example of a lipid-binding protein from *Bdellovibrio bacteriovorus* HD100 - the predation-associated TULIP, TipA, and identified an associated adapter protein TipB.

TipA is in the same superfamily as the eukaryotic bacterial permeability increasing protein (BPI), which also forms a double-TULIP fold. The protein shows similar structural features to that of other BPI-like proteins, such as the classical Helix-ββββ-Helix TULIP core secondary structure, two TULIP domains back-to-back with an intervening central sheet, and longer N-terminal than C-terminal TULIP domain. However, modifications to the protein have made the overall fold divergent, with the addition of molecular features that enable the formation of a dimeric double-TULIP, and for interaction with the TipB adapter protein. The dimerisation motif includes an additional hairpin and helix that lock the homodimer together to form a hugely elongated ∼260 Å tube.

While no other double-TULIP has been observed in a dimeric form, the extended synaptotagmins (E-Syts) and the eukaryotic ER-mitochondria encounter structure (ERMES) proteins do form single-TULIP homodimers, but they pack using a different interface where helix B packs against the beta sheet edge [39]. It is interesting that a heterotetrameric form of TULIP is also seen between ERMES proteins Mdm12 and Mmm1, where heterodimers stack head-to-head through an Mmm1-Mmm1 interface [40]. This creates an equivalent chain of four TULIP domains, highlighting that higher-order arrangements of TULIPs are functionally significant. It is possible that this increase in TULIP length is needed in order to efficiently transport lipids effectively at some juncture between prey and *Bdellovibrio*. The length of the dimer is between 250 and 260 Å which is equivalent to that of the periplasmic spanning transporter complexes, Yeb and Pqi [41] as well as the predicted structure of the putative transporter YhdP [42]. ERMES proteins and E-Syts tether membranes and transfer lipids between them, which is dependent on the presence of their C2 domains for membrane tethering [43], and therefore proximity of membranes to their single-TULIP is required. The increased length of the TipA dimer could allow for lipid transfer between membranes without the need for direct membrane tethering by additional domains.

It is also noteworthy that we consistently observe TipA packing C-terminal hydrophobic residues into the N-terminal domain – which creates a network of TULIPs in the crystal lattice. This could indicate that during predation TipA could form higher order structures that facilitate transport of lipids from TULIP to TULIP. Higher order structures also occur in the ERMES complex as the Mmm1-Mdm12 complex also interacts with another TULIP Mdm34 [40, 44], but structural information and exact stoichiometry is unknown for these larger complexes. Further work could confirm whether there is a physiological function for this consistent C-terminal-N-terminal TULIP packing of TipA.

The *tipA* gene is in an operon with *tipB*, which encodes for a small C2-like sandwich domain. The functional link between TULIPs and C2 domains is observed in ERMES and E-Syt systems, where the C2 domains mediate membrane tethering. TipB, while C2-like, does not bind calcium or membranes, and functions in an entirely different manner. The TipB sandwich has been adorned with two extended loops that are used for interaction with TipA by insertion into its lumen. The crystal structure of the complex of TipA and TipB shows that upon binding of the two proteins there is a large conformational change in TipA, mediated by a clamping mechanism that lifts its N-terminal beta sheet. This creates an opening of the TULIP seam, creating an accessible core across both N-terminal domains of the TipA dimer. Previously, a gate-and-latch mechanism has been proposed to account for lipid binding into the lumen of the single-TULIP juvenile-hormone binding protein from the domestic silk moth [45]. Here, the N-terminal helix A opens in the apo form and closes on the buried sesquiterpene molecule when bound. As the N-terminal helix of double-TULIP proteins is locked into the central beta sheet by an additional N-terminal beta strand, this latch mechanism is not possible. Instead, it is possible that the opening of TipA by TipB is what facilitates loading or unloading of lipids; observation of lipid-loaded recombinant TipA by TLC indicates that it would not be dependent on TipB. The intriguing finding of a planctomycete double-TULIP with a fused beta-sandwich that binds to the N-terminal domain Helix A and sheet proposes the idea that this functional relationship is repeated across diverse bacteria. In addition, it is possible that TipA homodimerisation is also required to ensure the stability of the overall fold during this opening and closing process as we have observed a high degree of flexibility in the N-terminal sheet (very pronounced in TipA_Δhairpin_), and the lumen of the N-terminal domain is larger than typically observed in other double-TULIPs.

We were unsuccessful in attempts to monitor TipA localisation – trials with a C-terminal mCherry tag resulted in frequent mutational escape, so TipA could not be investigated by fluorescence microscopy; future studies could use antibodies or computational-design binders to address this issue. TipA’s ability to adhere to membranes and destabilise them as shown via QCM-D suggests that it could be used to extract lipids directly from prey bacterial membranes. Binding and membrane destabilisation by TipA could aid *Bdellovibrio*, and could work in a similar fashion to BPI [46] by increasing membrane permeability. TipA is not the first example of a TULIP protein adhering to membranes without C2 domain partners - tricalbin 2 (a yeast E-syt homologue) and nucleus-vacuole junction protein 1 TULIP domains were shown to bind to membranes *in vitro* using liposomes [47], and the tricalbins are known to transfer lipids between membrane contact sites. Therefore, membrane adherence (as observed by both our purification procedure and QCM-D experiments) is likely an effective way for TipA to uptake lipids into its core ready for transfer to another protein or membrane.

In conclusion, our findings underscore the evolutionary significance of TULIPs as universal lipid-binding folds across both eukaryotes and prokaryotes. The crystal structures of TipA, TipB and their binary complex show that *B. bacteriovorus* has novel lipid-binding machinery for its predatory lifestyle in the form of a modified dimeric double-TULIP protein. Through the mechanistic interdependence of TipA and TipB, we give new insight into the conformational changes possible in double-TULIP proteins and suggest an adaptor-mediated mechanism for promoting lipid loading and unloading.

## Methods

### Cloning, expression and purification

TipA-His was cloned into a modified version of the pET41c overexpression vector using restriction-free cloning methods. The final construct contained residues of 26-572 of TipA (Uniprot accession Q6MK73) fused to a C-terminal octa-His-tag. TipB was cloned into pET26b using NdeI and NotI. A tagless form of TipA was made by mutagenesis to remove the octa-Histag. A monomeric form of tagless TipA (TipA_ΔHairpin_) was created using Q5 mutagenesis, replacing the dimer-hairpin residues A249-Y-258 (AVEVNGKKFY) with a triple glycine linker. TipB was cloned into pET26b using NdeI and XhoI restriction sites. The final construct contained an N-terminal MAHHHHHH tag and residues 29-171 of TipB (Uniprot accession Q6MK72).

To express selenomethionine-labelled TipA, a 60 mL overnight culture of BL21 (DE3) cells in LB was centrifuged and resuspended in minimal media supplemented with kanamycin and chloramphenicol to a final volume of 1 L. At OD_600_ 0.4, 0.1 g of lysine, 0.1 g of threonine, 0.1 g of phenylalanine, 0.05 g leucine, 0.05 g of isoleucine, 0.05 g of valine and 0.06 g of selenomethionine were added to the culture. After 15 minutes, IPTG was added to 1 mM. Cells were harvested by centrifugation after 16 h at 18 °C and stored at −20 °C.

TipA_ΔHairpin_, TipA-His and TipB constructs were expressed in *Escherichia coli* BL21 RIPL (DE3). Cells were grown in LB supplemented with kanamycin to an OD_600_ of 0.5-0.7 and induced with 0.5 mM IPTG at 18 °C for 16 h. Cells were harvested by centrifugation at 4000g for 15 minutes. During trial purifications we noted that the majority of TipA protein sedimented with insoluble cell components following lysis and centrifugation, but could be liberated from the insoluble pellet by the addition of high (1M) NaCl. We exploited this characteristic of TipA in our subsequent purification protocols. Cells expressing untagged selenomethionine-labelled TipA and TipA_ΔHairpin_ were resuspended in wash buffer (50 mM HEPES and 0.15 M NaCl, pH 7.5), sonicated and centrifuged at 48384 g for 1 h. The pellet was again resuspended, sonicated and centrifuged to wash away soluble proteins. A third pellet wash was performed and finally the pellet was resuspended in 50 mM HEPES and 1 M NaCl, pH 7.5. Cells expressing his-tagged proteins were resuspended in buffer A (TipA-His - 50 mM HEPES and 1M NaCl, pH 7.5; TipB - 50 mM HEPES and 300 mM NaCl, pH 7.5). The cells were lysed by sonication on ice and centrifuged at 48384 g for 1 hour. The pellet was resuspended. The lysates were loaded onto a 5 mL Ni-NTA column (GE Healthcare) and washed with 10 CV of lysis buffer. The protein was eluted with a gradient of 20–500 mM imidazole. All proteins were finally passed through a Superdex 200 pg or 75 pg 26/60 column (GE Healthcare), which was pre-equilibrated with 20 mM HEPES, 150 mM NaCl, pH 7.5. Fractions containing the protein of interest were pooled, concentrated and stored frozen at −70 °C if not required for immediate use.

### Crystallisation

Selenomethionine-lebelled TipA crystallisation screens were prepared by mixing a 1.5 μL of 8 mg/mL TipA and 1.5 μL precipitant solution. TipA crystallised in space group P 31 2 1 (a =252.4, b=252.4, c=181.1, α=90, β=90, γ=120) in a precipitant solution from the Jena Bioscience JBS Classic Screen consisting of 15% w/v PEG 4000; 100 mM trisosodium citrate, pH 5.6; 200 mM ammonium sulphate. Diffraction data were collected at Diamond Light Source. Data reduction and processing were performed using XDS.

TipA_ΔHairpin_ crystallisation screens were prepared by mixing a 0.4 μL of 10 mg/mL TipA and 0.4 μL precipitant solution. TipA crystallised in space group P 21 21 21 (a =58.0, b=97.8, c=132.0, α=90, β=90, γ=90) in a precipitant solution from the BCS screen consisting of 0.1 M MES pH 6.5, 2% v/v PEG 400, 2% v/v PEG 500 MME, 2% v/v PEG 600, 2% w/v PEG 1000, 2% w/v PEG 2000, 2% w/v PEG 3350, 2% w/v PEG 4000, 2% w/v PEG 5000 MME, 2% w/v PEG 6000, 2% w/v PEG 8000, 2% w/v PEG 10000. Diffraction data were collected at Diamond Light Source. Data reduction and processing were performed through the Xia-2 Dials pipline, and Aimless.

TipB crystallisation screens were prepared by mixing a 0.4 μL of 10 mg/mL TipB and 0.4 μL precipitant solution. TipB crystallised in space group C 2 2 21 (a =106.6, b=126.2, c=78.0, α=90, β=90, γ=90) in a precipitant solution from the Morpheus screen consisting of 0.03 M Diethylene glycol; 0.03 M Triethylene glycol; 0.03 M Tetraethylene glycol; 0.03 M Pentaethylene glycol, 0.1 M Bicine, 0.1 M Tris, pH 8.5, 20% v/v PEG 500 MME, 10 % PEG 20000. Diffraction data were collected at Diamond Light Source. Data reduction and processing were performed through the Autoproc suite, and Aimless.

TipAB crystallisation screens were prepared by mixing a 0.5 μL of 8 mg/mL TipA and 9.5 mg/mL TipB (approx. 5X molar ratio) and 0.5 μL precipitant solution. The TipAB complex crystallised in space group C 2 2 21 (a =89.3, b=230.7, c=131.6, α=90, β=90, γ=90) in a precipitant solution from the Proplex screen consisting of 0.1 M Sodium acetate pH 5, 0 2 % w/v PEG 4000 15 % v/v MPD. Diffraction data were collected at European Synchrotron Radiation Facility. Data reduction and processing were performed through the Xia2-Dials pipeline.

### Structure determination and model building

Diffraction data processing and refinement statistics are given in Table EV1.

The crystal structure of Selenomethionine-labelled TipA was solved using Phenix, with manual interpretation and improvement of non-crystallographic symmetry operators.

The crystal structure of TipB was solved by molecular replacement with a Colabfold model [48] in Phenix MR [49] Likewise, TipA_Δhairpin_ was solved by molecular replacement with the C-terminal domain of the selenomethionine-labelled TipA and manual boot-strap building and refinement of the N-terminal domain. The complex TipAB structure was solved using molecular replacement of the TipB structure with the interaction loops removed, and the C-terminal domain of the selenomethionine-labelled TipA structure. The N-terminal domain was also manually built through boot-strap building and refinement. All structures were refined through iterative cycles of real-space building in Coot [50] and refinement in Phenix Refine [51].

### Pellet binding assays

Untagged TipA was expressed in *Escherichia coli* BL21 RIPL (DE3). Cells were grown in 10 mL LB supplemented with kanamycin to an OD_600_ of 0.5-0.7 and induced with 0.5 mM IPTG at 18 °C for 16 h. 5 × 1 mL of cells were centrifuged at 13000 RPM for 15 seconds and the supernatant was removed. The cells were resuspended in 500 μL of either 0.15 M NaCl, 0.05 M HEPES pH 7.5; 0.5 M NaCl, 0.05 M HEPES pH 7.5; 1 M NaCl, 0.05 M HEPES pH 7.5; 0.15 M NaCl, 0.05 M HEPES pH 7.5, 0.05% Triton X100; 0.15 M NaCl, 0.05 M HEPES pH 7.5, 0.05% Tween 20. sonicated for 15 seconds and centrifuged at 13000 RPM for 15 minutes. The pellet obtained from the 0.15 M NaCl buffer was resuspended in 500 μL SDS-PAGE loading buffer for the pellet control. The pellet control and supernatant of each buffer was run on SDS-PAGE.

### QCM-D

QCM-D was performed using a Q-sense Analyzer QCM-D System (Biolin Scientific, Gothenburg, Sweden). Silicon QCM-D sensors were fixed into the QCM-D flow cell at a constant temperature of 20 °C and attached to a peristaltic pump. ddH_2_O was pumped through the flow cell at a constant rate of 0.1 ml/min, a rate which was subsequently maintained throughout the rest of the experiment. Frequency and dissipation changes (Δf and ΔD) were monitored using multiple harmonics (n = 3, 5, 7, 9, 11, 13) of the resonant frequency. For simplicity we report here only the changes to the 3rd harmonic. Prior to data acquisition flow cells were equilibrated in H_2_O for 5 mins to allow for a stable baseline to be acquired. H2O was then exchanged for buffer, which was pumped through the flow cell until the change in frequency of the 3rd harmonic was stabilised. Bilayers consisting of either POPC, *E. coli* polar lipids or POPC:LPS (LPS from *E. coli* O111:B4) were then deposited onto the silicon sensor. This initially involves the formation of small unilamellar vesicles (SUVs), which were prepared by resuspension of a desiccated POPC/E. coli Polar Lipids/POPC:LPS film in buffer in 150 mM NaCl, 20 mM HEPES pH 7.5 to a concentration of 0.2 mg/mL and subsequent sonication for 30 min until an optically transparent solution was obtained. POPC:LPS SUVs were prepared at a 10:1 (w/w) ratio. These SUVs were subsequently flowed over the silicon sensor and then ruptured through osmotic shock with H2O, resulting in the characteristic formation of a deposited lipid bilayer. Exchange back into buffer 150 mM NaCl, 20 mM HEPES pH 7.5 aids the formation of a stable bilayer, with Δf3 ranging between −35 to −45 Hz. Once bilayers were deposited and baselines stabilized, TipA was introduced into the flow cell at a concentration of 1 mg/ml and flowed over the deposited bilayers for ∼45 mins. Excess TipA was then removed by washing sensors in 150 mM NaCl, 20 mM HEPES pH 7.5. QCM-D with TipB was performed as above for TipA with POPC bilayers.

An *E. coli* Polar Lipid:LPS (10:1 w/w) mixed bilayer was also attempted to be formed and deposited onto silicon surfaces as outlined above. These bilayers were unable to be successfully generated, with SUVs unable to rupture and subsequently washed from the sensor. These surfaces were treated as ‘unformed bilayers’ and used as control surfaces to monitor TipA interaction with empty silicon sensors.

### ITC

To test whether TipB binds to calcium, 5 mM EDTA and 5 mM EGTA was added to the protein following size exclusion. After 16 hours the protein was passed through a Superdex 75 pg 26/60 column (GE Healthcare), which was pre-equilibrated with 20 mM HEPES, 150 mM dialysed against 1 L of 20 mM HEPES, 150 mM NaCl, pH 7.5. CaCl_2_ was dissolved in the dialysis buffer to a final concentration of 500 μM. ITC was performed by titrating 500 μM CaCl_2_ into 50 μM TipB in 13 injections (1 x 0.4, 12 x 3 μL injections).

To test whether TipB binds to TipA-His, purified protein was dialysed against 1 L of 20 mM HEPES, 150 mM NaCl, pH 7.5. ITC was performed by titrating 200 μM TipB into 20 μM TipA in 13 injections (1 x 0.4, 12 x 3 μL injections).

### TLC

TLC was employed to assess the lipid binding ability of TipA. Lipids were extracted from protein samples through chloroform/methanol extraction. Briefly, 2 ml of Bd2538 at 0.5 mg/ml was added to 2 ml methanol and 1 ml chloroform (2:2:1, v/v/v). Samples were vortexed for 5 min, followed by a 30 min incubation at 50 °C and then vortexed again for 5 min. Centrifugation at 2,000 x g for 10 min resulted in the formation of two phases, the lower chloroform layer containing lipidic material. This lower phase was extracted and then evaporated. The dried lipids were resuspended in 100 μl chloroform with 5 μl then loaded onto a Silica TLC plate (Sigma) and run in a 6.5:2.5:1 (chloroform:methanol:acetic acid) solvent system. The TLC plate was dried, stained with 10% (w/v) phosphomolybdic acid in ethanol, and finally heated until stained spots could be visualised.

---

**Figure EV1.**
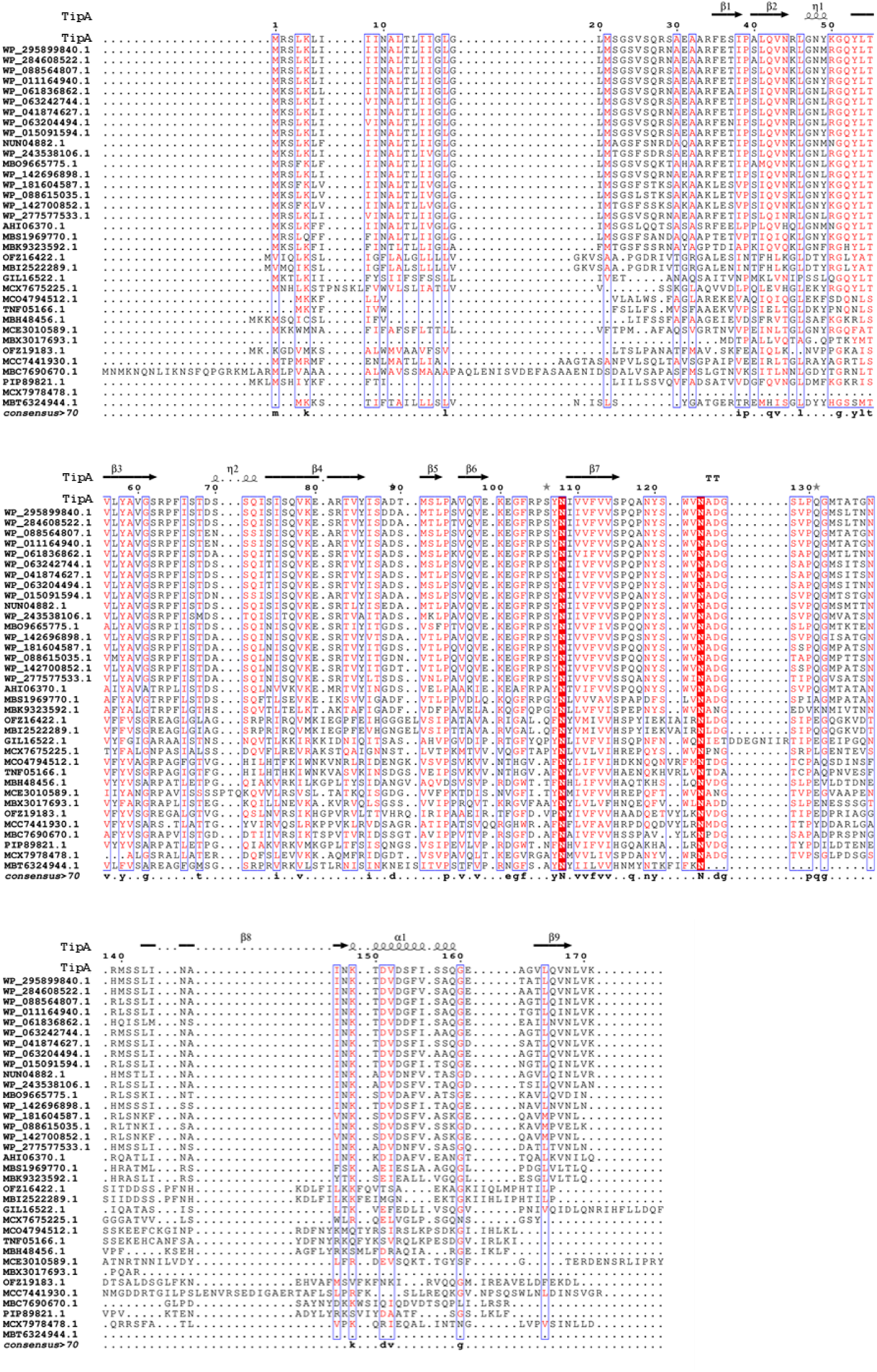
Sequence alignments of TipA homologues. BlastP [55] was used to identify homologues to the *Bdellovibrio bacteriovorus* HD100 TipA protein. The only absolutely conserved residues are two Asn residiues. Alignments of the sequences obtained from blastp were made using Clustal Omega [56]. ESPRIPT [57] was used to visualise the alignement with secondary structure.

**Figure EV2.**
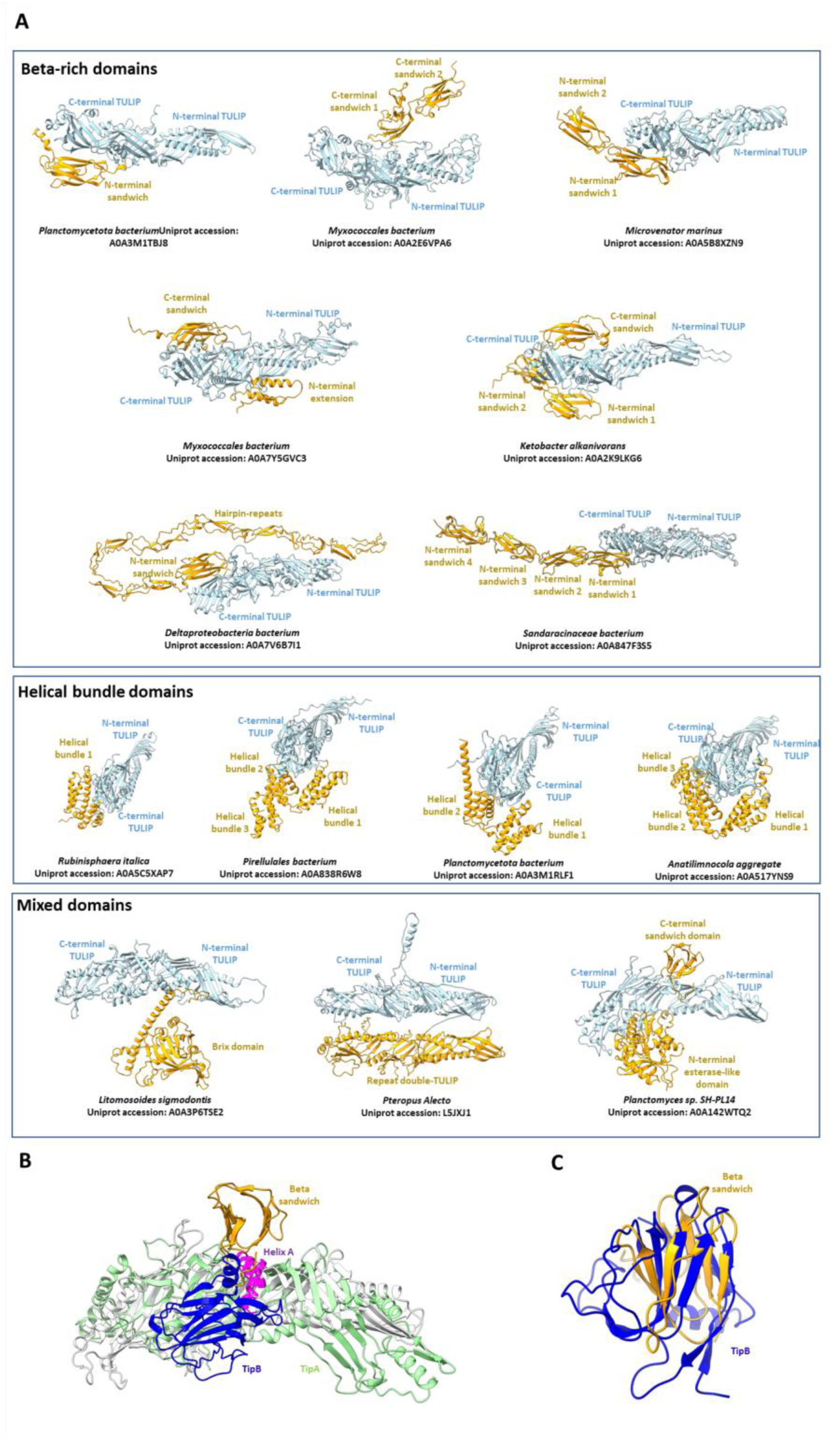
Additional domains fused to TipA structural homologues. **(A)** Using Foldseek [38], structural TipA homologues were identified within the AlphaFold [58] AFDB50 database. A number of those homologues had additional domains to the double-TULIP. Representative examples of fusions are shown and comprise beta-rich domains (upper) helical domains (middle) and other domains including second double-TULIPs and enzymes (lower). The double-TULIP domains are coloured light blue, and the accessory domains are coloured orange. Of particular interest is the *Planctomyces sp*. SH-PL14 protein (bottom right) in which there is a similar beta-sandwich domain to TipB fused C-terminally to the TULIP **(B)** Superposition of the TipAB with *Planctomyces sp*. SH-PL14 (Uniprot accession: A0A142WTQ2). TipA is coloured green, TipB is blue, the TULIP domains of A0A142WTQ2 are white and the beta sandwich is orange. The enzymatic domain has been removed for clarity. The beta-sandwiches interact with the N-terminal Helix A in both structures and make interactions with the top of the N-terminal TULIP sheet. **(C)** Superposition of TipB (blue) with the beta sandwich of A0A142WTQ2 (orange), showing similar topology.

**Figure EV3.**
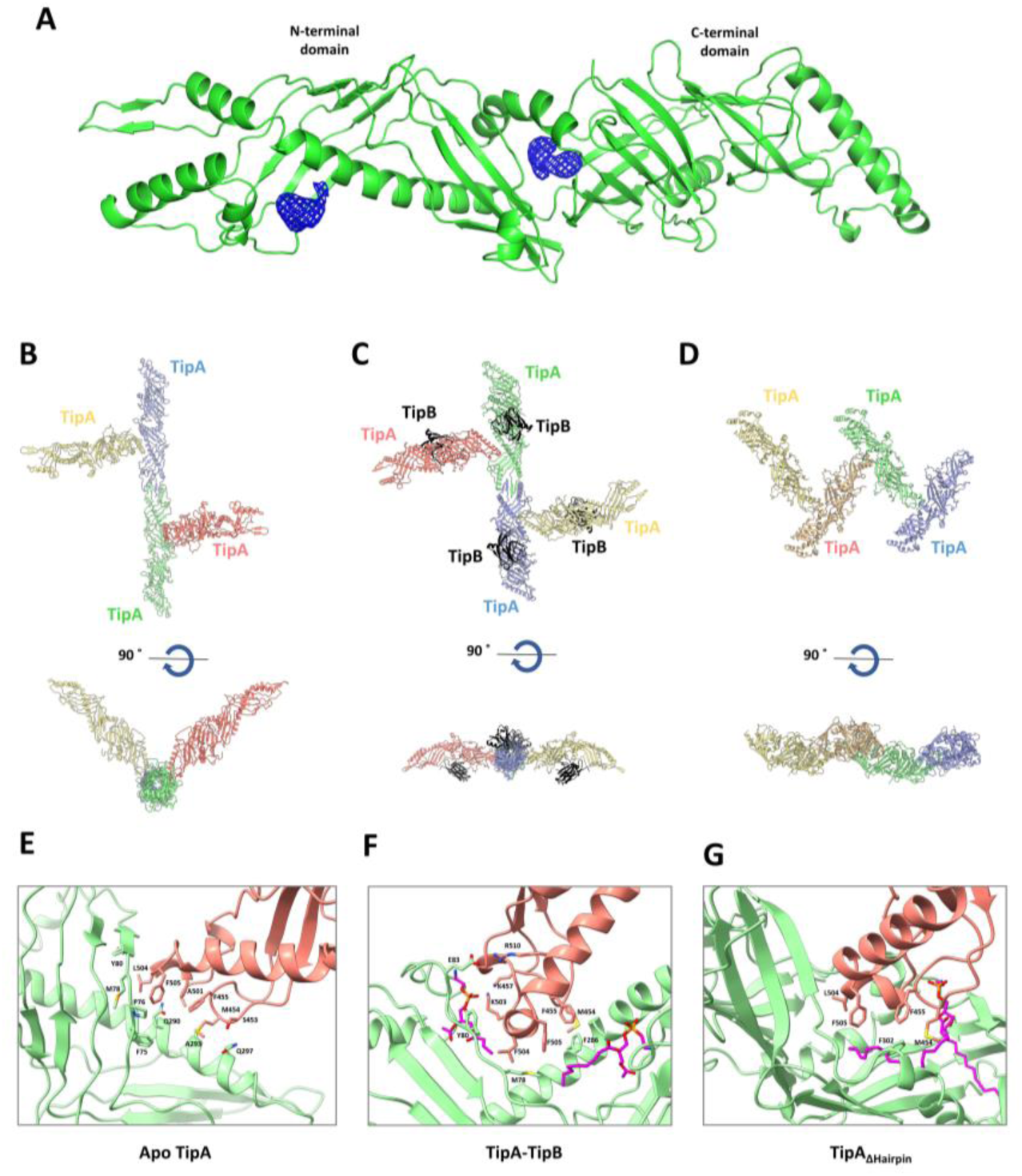
Lipid binding and higher order packing of TipA. **(A)** Fo-Fc electron density (blue mesh) observed in the apo selenomethionine TipA structure. Large peaks are observed in similar regions to lipid density in the TipAB complex and TipA_ΔHairpin_. **(B)** Crystal packing in the apo selenomethionine structure. **(C)** Crystal packing in the TipAB complex structure. **(D)** Crystal packing in the TipA_ΔHairpin_ structure. In all structures shown **(B-D)** the C-terminal domain of TipA buries into the N-terminal domain of another TipA protein. **(E)** Molecular interactions of the C-terminal domain and N-terminal of the apo selenomethionine TipA structure. **(F)** Molecular interactions of the C-terminal domain and N-terminal of the TipAB structure. **(G)** Molecular interactions of the C-terminal domain and N-terminal of the TipA_ΔHairpin_ structure.

**Table EV1.**
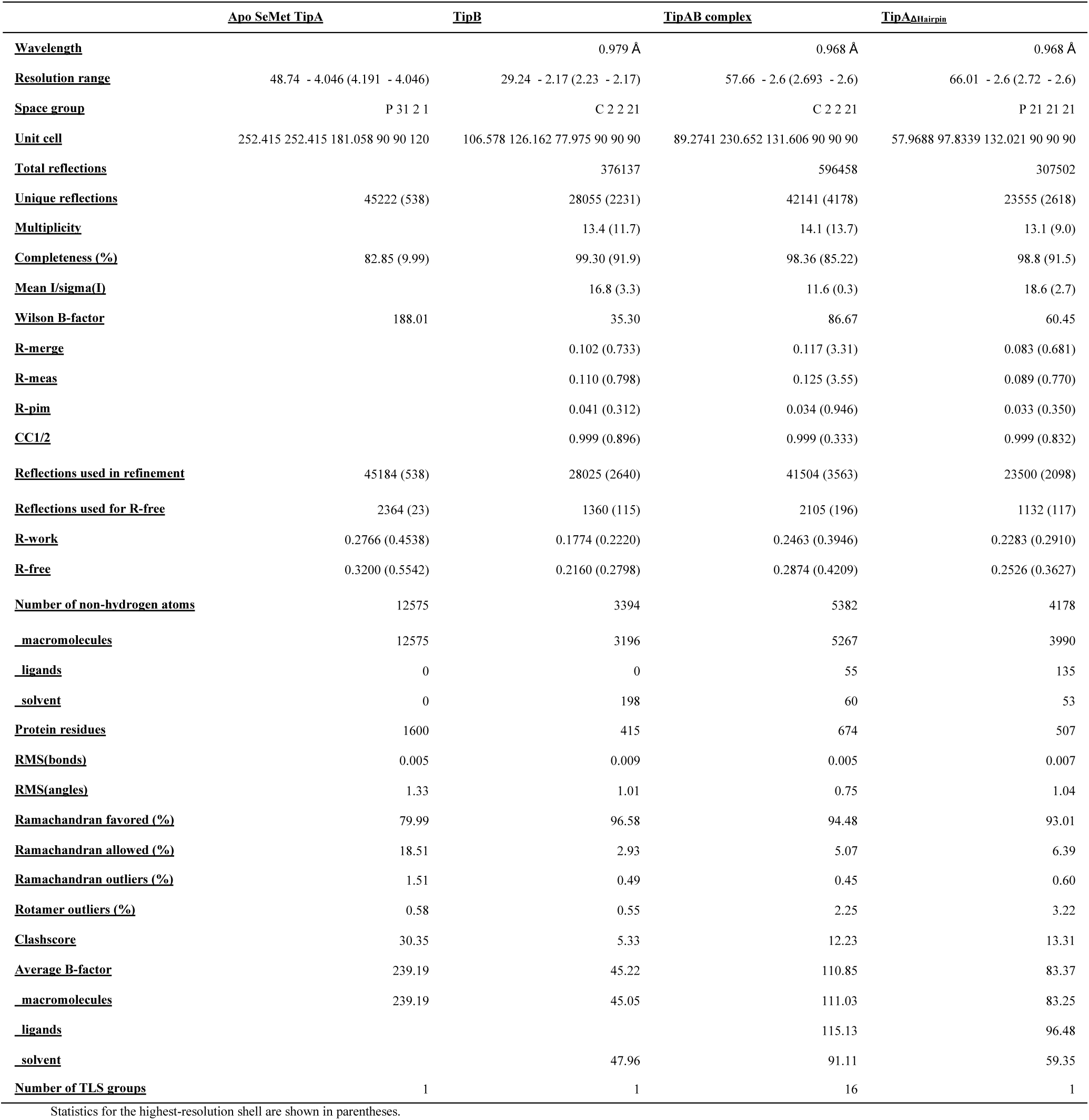
Data collection and refinement statistics.

## Acronyms

ASU: Asymmetric unit
TULIP: Tubular lipid binding protein
SMP: Synaptotagmin-like mitochondrial-lipid-binding protein
LPT: Lipopolysaccharide transport
MLA: Maintenance of OM lipid asymmetry
BPI: Bacterial permeability-increasing protein
CETP: Cholesteryl ester transfer protein
LBP: LPS-binding protein
E-Syt: Extended Syntaptotagmin
ERMES: ER mitochondrial encounter structure
RMSD: Root mean square deviation
TLC: Thin layer chromatography
ITC: Isothermal titration calorimetry
QCM-D: Quartz crystal microbalance with dissipation
PDB: protein data bank
CL: cardiolipin
PE: Phosphatidylethanolamine
PG: phosphatidylglycerol
AFDB50: AlphaFold Database 50

## Acknowledgements

ALL, SGC, and ITC were funded by the Wellcome Trust Investigator Award in Science (209437/Z/17/Z); AL and SGC acknowledge additional funding from a BBSRC award (BB/X006298/1). GH and TJK were funded by BBSRC award BB/S017283/1, and HJ by BBSRC studentship BB/T00746X/1. We are grateful to Paul Radford and Liz Sockett for attempts to monitor functional role and localization in vivo. We thank Diamond Light Source for beamtime and for facility support during data collection.

## References

1. Negus, D., et al., Predator Versus Pathogen: How Does Predatory Bdellovibrio bacteriovorus Interface with the Challenges of Killing Gram-Negative Pathogens in a Host Setting? Annu Rev Microbiol, 2017. 71: p. 441–457.

2. Bratanis, E., et al., Biotechnological Potential of Bdellovibrio and Like Organisms and Their Secreted Enzymes. Front Microbiol, 2020. 11: p. 662.

3. Rendulic, S., et al., A predator unmasked: life cycle of Bdellovibrio bacteriovorus from a genomic perspective. Science, 2004. 303(5658): p. 689–92.

4. Matin, A. and S.C. Rittenberg, Kinetics of deoxyribonucleic acid destruction and synthesis during growth of Bdellovibrio bacteriovorus strain 109D on pseudomonas putida and escherichia coli. J Bacteriol, 1972. 111(3): p. 664–73.

5. Hespell, R.B., G.F. Miozzari, and S.C. Rittenberg, Ribonucleic acid destruction and synthesis during intraperiplasmic growth of Bdellovibrio bacteriovorus. J Bacteriol, 1975. 123(2): p. 481–91.

6. Thomashow, M.F. and S.C. Rittenberg, Intraperiplasmic growth of Bdellovibrio bacteriovorus 109J: solubilization of Escherichia coli peptidoglycan. J Bacteriol, 1978. 135(3): p. 998–1007.

7. Rosson, R.A. and S.C. Rittenberg, Regulated breakdown of Escherichia coli deoxyribonucleic acid during intraperiplasmic growth of Bdellovibrio bacteriovorus 109J. J Bacteriol, 1979. 140(2): p. 620–33.

8. Nelson, D.R. and S.C. Rittenberg, Incorporation of substrate cell lipid A components into the lipopolysaccharide of intraperiplasmically grown Bdellovibrio bacteriovorus. J Bacteriol, 1981. 147(3): p. 860–8.

9. Kuenen, J.G. and S.C. Rittenberg, Incorporation of long-chain fatty acids of the substrate organism by Bdellovibrio bacteriovorus during intraperiplasmic growth. J Bacteriol, 1975. 121(3): p. 1145–57.

10. Abram, D., J. Castro e Melo, and D. Chou, Penetration of Bdellovibrio bacteriovorus into host cells. J Bacteriol, 1974. 118(2): p. 663–80.

11. Banks, E.J., et al., Asymmetric peptidoglycan editing generates cell curvature in Bdellovibrio predatory bacteria. Nat Commun, 2022. 13(1): p. 1509.

12. Kuru, E., et al., Fluorescent D-amino-acids reveal bi-cellular cell wall modifications important for Bdellovibrio bacteriovorus predation. Nat Microbiol, 2017. 2(12): p. 1648–1657.

13. Giacometti, S.I., et al., Lipid Transport Across Bacterial Membranes. Annu Rev Cell Dev Biol, 2022. 38: p. 125–153.

14. Reinisch, K.M. and P. De Camilli, SMP-domain proteins at membrane contact sites: Structure and function. Biochim Biophys Acta, 2016. 1861(8 Pt B): p. 924–927.

15. Alva, V. and A.N. Lupas, The TULIP superfamily of eukaryotic lipid-binding proteins as a mediator of lipid sensing and transport. Biochim Biophys Acta, 2016. 1861(8 Pt B): p. 913–923.

16. Wong, L.H. and T.P. Levine, Tubular lipid binding proteins (TULIPs) growing everywhere. Biochim Biophys Acta Mol Cell Res, 2017. 1864(9): p. 1439–1449.

17. Levine, T.P., Remote homology searches identify bacterial homologues of eukaryotic lipid transfer proteins, including Chorein-N domains in TamB and AsmA and Mdm31p. BMC Mol Cell Biol, 2019. 20(1): p. 43.

18. Barrio-Hernandez, I., et al., Clustering predicted structures at the scale of the known protein universe. Nature, 2023. 622(7983): p. 637–645.

19. Gustafsson, R., et al., Crystal structures of OrfX2 and P47 from a Botulinum neurotoxin OrfX-type gene cluster. FEBS Lett, 2017. 591(22): p. 3781–3792.

20. Gao, L., et al., Crystal structures of OrfX1, OrfX2 and the OrfX1-OrfX3 complex from the orfX gene cluster of botulinum neurotoxin E1. FEBS Lett, 2023. 597(4): p. 524–537.

21. Lambert, C., et al., The first bite--profiling the predatosome in the bacterial pathogen Bdellovibrio. PLoS One, 2010. 5(1): p. e8599.

22. Tyson, J., et al., Prey killing without invasion by Bdellovibrio bacteriovorus defective for a MIDAS-family adhesin. Nat Commun, 2024. 15(1): p. 3078.

23. Paysan-Lafosse, T., et al., InterPro in 2022. Nucleic Acids Res, 2023. 51(D1): p. D418–D427.

24. Holm, L., Dali server: structural unification of protein families. Nucleic Acids Res, 2022. 50(W1): p. W210–W215.

25. Voss, N.R. and M. Gerstein, 3V: cavity, channel and cleft volume calculator and extractor. Nucleic Acids Res, 2010. 38(Web Server issue): p. W555–62.

26. Izore, T., et al., Structural characterization and membrane localization of ExsB from the type III secretion system (T3SS) of Pseudomonas aeruginosa. J Mol Biol, 2011. 413(1): p. 236–46.

27. Nalefski, E.A. and J.J. Falke, The C2 domain calcium-binding motif: structural and functional diversity. Protein Sci, 1996. 5(12): p. 2375–90.

28. Kohout, S.C., et al., C2 domains of protein kinase C isoforms alpha, beta, and gamma: activation parameters and calcium stoichiometries of the membrane-bound state. Biochemistry, 2002. 41(38): p. 11411–24.

29. Nalefski, E.A., et al., Delineation of two functionally distinct domains of cytosolic phospholipase A2, a regulatory Ca(2+)-dependent lipid-binding domain and a Ca(2+)-independent catalytic domain. J Biol Chem, 1994. 269(27): p. 18239–49.

30. Saheki, Y. and P. De Camilli, The Extended-Synaptotagmins. Biochim Biophys Acta Mol Cell Res, 2017. 1864(9): p. 1490–1493.

31. Dixon, M.C., Quartz crystal microbalance with dissipation monitoring: enabling real-time characterization of biological materials and their interactions. J Biomol Tech, 2008. 19(3): p. 151–8.

32. Hughes, G.W., et al., Evidence for phospholipid export from the bacterial inner membrane by the Mla ABC transport system. Nat Microbiol, 2019. 4(10): p. 1692–1705.

33. Richter, R., A. Mukhopadhyay, and A. Brisson, Pathways of lipid vesicle deposition on solid surfaces: a combined QCM-D and AFM study. Biophys J, 2003. 85(5): p. 3035–47.

34. Shen, H.H., et al., Reconstitution of a nanomachine driving the assembly of proteins into bacterial outer membranes. Nat Commun, 2014. 5: p. 5078.

35. Perisic, O., et al., Crystal structure of a calcium-phospholipid binding domain from cytosolic phospholipase A2. J Biol Chem, 1998. 273(3): p. 1596–604.

36. Krissinel, E. and K. Henrick, Inference of macromolecular assemblies from crystalline state. J Mol Biol, 2007. 372(3): p. 774–97.

37. Lee, R.A., M. Razaz, and S. Hayward, The DynDom database of protein domain motions. Bioinformatics, 2003. 19(10): p. 1290–1.

38. van Kempen, M., et al., Fast and accurate protein structure search with Foldseek. Nat Biotechnol, 2024. 42(2): p. 243–246.

39. Schauder, C.M., et al., Structure of a lipid-bound extended synaptotagmin indicates a role in lipid transfer. Nature, 2014. 510(7506): p. 552–5.

40. AhYoung, A.P., et al., Conserved SMP domains of the ERMES complex bind phospholipids and mediate tether assembly. Proc Natl Acad Sci U S A, 2015. 112(25): p. E3179–88.

41. Ekiert, D.C., et al., Architectures of Lipid Transport Systems for the Bacterial Outer Membrane. Cell, 2017. 169(2): p. 273–285 e17.

42. Kumar, S. and N. Ruiz, Bacterial AsmA-Like Proteins: Bridging the Gap in Intermembrane Phospholipid Transport. Contact (Thousand Oaks), 2023. 6: p. 25152564231185931.

43. Yu, H., et al., Extended synaptotagmins are Ca2+-dependent lipid transfer proteins at membrane contact sites. Proc Natl Acad Sci U S A, 2016. 113(16): p. 4362–7.

44. Jeong, H., J. Park, and C. Lee, Crystal structure of Mdm12 reveals the architecture and dynamic organization of the ERMES complex. EMBO Rep, 2016. 17(12): p. 1857–1871.

45. Suzuki, R., et al., Structural mechanism of JH delivery in hemolymph by JHBP of silkworm, Bombyx mori. Sci Rep, 2011. 1: p. 133.

46. Weiss, J., et al., Purification and characterization of a potent bactericidal and membrane active protein from the granules of human polymorphonuclear leukocytes. J Biol Chem, 1978. 253(8): p. 2664–72.

47. Toulmay, A. and W.A. Prinz, A conserved membrane-binding domain targets proteins to organelle contact sites. J Cell Sci, 2012. 125(Pt 1): p. 49–58.

48. Mirdita, M., et al., ColabFold: making protein folding accessible to all. Nat Methods, 2022. 19(6): p. 679–682.

49. McCoy, A.J., Solving structures of protein complexes by molecular replacement with Phaser. Acta Crystallogr D Biol Crystallogr, 2007. 63(Pt 1): p. 32–41.

50. Emsley, P. and K. Cowtan, Coot: model-building tools for molecular graphics. Acta Crystallogr D Biol Crystallogr, 2004. 60(Pt 12 Pt 1): p. 2126–32.

51. Afonine, P.V., et al., Towards automated crystallographic structure refinement with phenix.refine. Acta Crystallogr D Biol Crystallogr, 2012. 68(Pt 4): p. 352–67.

52. Sehnal, D., et al., MOLE 2.0: advanced approach for analysis of biomacromolecular channels. J Cheminform, 2013. 5(1): p. 39.

53. Hall, M., et al., Structural and functional characterization of shaft, anchor, and tip proteins of the Mfa1 fimbria from the periodontal pathogen Porphyromonas gingivalis. Sci Rep, 2018. 8(1): p. 1793.

54. Adhikari, B., et al., DISTEVAL: a web server for evaluating predicted protein distances. BMC Bioinformatics, 2021. 22(1): p. 8.

55. Altschul, S.F., et al., Basic local alignment search tool. J Mol Biol, 1990. 215(3): p. 403–10.

56. Sievers, F. and D.G. Higgins, Clustal Omega for making accurate alignments of many protein sequences. Protein Sci, 2018. 27(1): p. 135–145.

57. Robert, X. and P. Gouet, Deciphering key features in protein structures with the new ENDscript server. Nucleic Acids Res, 2014. 42(Web Server issue): p. W320–4.

58. Jumper, J., et al., Highly accurate protein structure prediction with AlphaFold. Nature, 2021. 596(7873): p. 583–589.

